# Hummingbird torpor recapitulates key molecular signatures of hibernation without large-scale transcriptomic remodelling

**DOI:** 10.64898/2026.09.27.753529

**Authors:** Harsha Kumar, Emily R Blackwell, Donald R Powers, Anusha Shankar

## Abstract

Torpor is a physiological strategy by which some species conserve energy by lowering metabolic rates and body temperatures. While mammalian hibernation is well-studied, the molecular mechanisms governing the rapid entry into avian torpor remain poorly understood. This study represents the first multi-tissue whole-transcriptomic analysis of torpor entry in an avian system. By combining real-time thermal imaging and respirometry with transcriptomic profiling across seven tissues, we examined the key molecular pathways involved in this transition in Anna’s hummingbirds (*Calypte anna*). Our results reveal that avian torpor entry is characterised by a subtle transcriptomic change (<5% of the genes tested differentially expressed across tissues) compared to mammalian hibernation (∼2–35%). About half of the genes whose expression responded to metabolic state or temperature were better described by physiological state than by surface temperature, whereas a smaller set tracked surface temperature continuously. The majority of the genes did not change expression between normothermy and transition. Vital organs, such as the heart and lungs, remained transcriptionally stable to maintain essential functions, while the liver and gut show the most change. We found no evidence of global transcriptional or translational suppression at the mRNA level. Instead, changes were selective and involved downregulation of splicing regulators and their kinases, upregulation of circadian repressors that are central to metabolism (NR1D1, BHLHE40), an upregulation of SGK1 and FOXO3, and slight downregulation in cell cycle and repair genes, consistent with a shift from growth towards cell maintenance and protection. Finally, the counterintuitive upregulation of genes involved in mitochondrial metabolism may reflect a preparation for rapid arousal despite the depressed metabolic state.

**Significance Statement:** Torpor allows animals to dramatically reduce energy expenditure during periods of energetic deficit, yet its molecular regulation in non-mammals remains poorly understood. Using multi-tissue transcriptomics, we examined the transition from normothermy to deep torpor in Anna’s hummingbirds. Interestingly, tissues undergo limited transcriptomic remodelling (1–4% of genes differentially expressed per tissue), unlike mammalian hibernators (∼15–60%). Hummingbirds regulate key pathways shared with mammals—lowering splicing regulators, raising clock repressors and cell-survival kinases, and priming mitochondrial metabolism for rapid arousal—without globally shutting down transcription or translation. The degree of transcriptomic changes observed suggests that avian torpor may rely more heavily on post-transcriptional and post-translational regulation than previously appreciated and provides a molecular framework for understanding the evolution of dormancy across vertebrates.

## Introduction

In the face of environmental pressures, such as food scarcity or low temperatures, some endothermic species use heterothermy by lowering their body temperatures and metabolic rates to save energy (1, 2). While seasonal hibernation involves multi-day bouts of deep metabolic suppression, daily torpor is a short-term state typically lasting less than a day (1–3). Although the molecular biology of mammalian hibernation and daily torpor has been extensively studied, avian torpor remains largely unexplored, with only studies on endocrine regulation (4–6), mitochondrial activity (7) and AKT-mTOR pathway regulation (8). Investigating the molecular basis of these dormant states can help determine whether the same pathways or molecular players regulate heterothermy or if these strategies have evolved independently across diverse taxonomic groups.

Mammalian hibernation, and, to a lesser extent, daily torpor, have been characterised at the molecular level, revealing highly coordinated transcriptomic reprogramming (3, 9). For instance, less than 5% of the heart transcriptome is differentially expressed in daily torpor in Djungarian hamsters (*Phodopus sungorous*), primarily affecting RNA stability, translational control, and oxygen handling (10). Global transcriptional suppression in mammalian torpor is accompanied with RNA-binding proteins (CIRBPs) that maintain transcript stability, and splicing factors (SRSFs) that alter splicing landscapes in ground squirrels (3, 9). Master regulators like FOXO3a induce cell cycle arrest (p27) and anti-oxidant responses (SOD2), while peripheral clocks (BMAL1, REV-ERBα) drastically modulate their amplitude during daily torpor or de-synchronise across tissue types in hibernation (10, 11). A hypothetical model proposes that torpor entry is initiated by fasting-associated increases in AMP and NAD^+^, which activate AMPK and sirtuin signalling to suppress glycolysis, enhance lipid utilisation, and trigger circadian-mediated metabolic reprogramming (12). This upstream signalling drives the characteristic fuel switch from carbohydrates to lipids via PPARα activation and PDK4 inhibition of carbohydrate oxidation (13). This switch to lipid utilisation is uniquely characterised by the upregulation of mitochondrial complex II, TCA cycle enzymes, and SOD2 defences in daily torpor in hamsters (14). While this upregulation was proposed to sustain elevated torpid body temperatures above ambient levels (14), the precise kinetic purpose of this mitochondrial priming remains open to interpretation. Cell lines of hibernating mammals when exposed to cold stress have shown an upregulation of anti-ferroptosis pathways (preventing cell death in the cold), with GPX4 playing a key role (15–17).

The molecular underpinnings of avian heterothermy have been poorly explored despite deep torpor representing one of the most extreme physiological shifts observed in vertebrates, characterised by sharp declines in heart rate, breathing rate, and whole-animal metabolic rate (1, 2). Endocrine signals like corticosterone (5) and seasonal fat thresholds (4, 18) have been shown to modulate torpor entry, while leptin lacks this regulatory role in hummingbirds (6) despite its known regulatory role in mammalian hibernation (19, 20). Deep torpor in hummingbirds can involve up to a 95% reduction in metabolic rate (2) driven by reduced ATP demands rather than an intrinsic downregulation of mitochondrial oxidative capacity, which might act as an adaptive feature for rapid arousal (7). The sole existing avian dataset on targeted torpor transcriptomics suggests tissue-specific re-weighting of Akt–mTOR signalling pathways, which conserves energy in skeletal muscles while preserving essential metabolic coordination and supporting anti-apoptotic functions in the liver and brain of speckled mousebirds (*Colius striatus*) (8). Which of these responses initiate torpor and which others follow from it, depends on when in torpor the tissues are sampled.

Different timepoints in the torpid state (early vs late torpor) represent distinct molecular states. For instance, hibernating greater horseshoe bats (*Rhinolophus ferrumequinum*) and thirteen-lined ground squirrels (*Ictidomys tridecemlineatus*) transition from broad transcriptional down-regulation in early torpor to selective gene expression or transcript stabilization in later torpor (21, 22). Because transcription effectively ceases at very low body temperatures in hibernation due to the shutdown of RNA Polymerase II kinetics, the increased relative abundance of certain transcripts during torpor suggests that these mRNAs are selectively stabilised rather than newly synthesised (23). Furthermore, these transitions are fundamentally constrained by the kinetics of gene expression. Although RNA Polymerase II transcribes at typical rates of 1 to 4 kilobases per minute (kb/min), splicing and processing most transcripts takes 5 to 30 minutes (24, 25). While sampling later into torpor would allow us to assess the genes involved in maintaining hypometabolism and transcript stability necessary for later arousal, the gene expression initiating torpor entry may be swamped by potential downstream secondary/compensatory programs by then. Focusing on early torpor therefore provides a critical window to identify the master regulators that initiate metabolic suppression and coordinate the initial molecular response downstream of hormonal and neuronal regulation.

In this study, we investigate the molecular signatures of early torpor in the Anna’s hummingbird (*Calypte anna*) to pinpoint the specific genes and pathways that orchestrate a rapid physiological transition from normothermy to torpor (Figure 1A). Daily torpor represents a different time scale and depth than hibernation, and heat loss kinetics in a small bird are likely very different from those of a large mammal. It is therefore interesting to concordantly examine the magnitude and rate of change in metabolic rates, body temperatures, and gene expression during torpor entry in these birds relative to daily torpid and hibernating mammals. Metabolic rates and body temperatures themselves have not been measured concurrently in birds in torpor, but evidence from bats demonstrates that these variables can become dynamically decoupled, allowing them to maintain profoundly suppressed metabolic rates even at high body temperatures (26, 27). Gene expression can either act as an upstream driver for metabolic suppression/change in body temperature or occur as a downstream consequence of these processes (14, 22, 28). Therefore, capturing the changes in transcriptomes of birds along the continuum of metabolic depression should illustrate the mechanistics, degree, and identity of players involved in torpor entry. We hypothesise that torpor entry in this system is regulated at the transcriptomic level and will be characterised by large-scale transcriptomic remodelling across tissue types (**Hypothesis 1**). Specifically, if the transition state is transcriptomically intermediate when compared to normothermy and torpor, this would be indicative of Q10 effects, while if it is similar in transition to either of the other two states, it would indicate active remodelling (**Hypothesis 2**). Given that the energy budgets of cells are dominated by the costs of translation, transcription and DNA replication (29, 30), we hypothesise that metabolic suppression in torpor will be coordinated by a suppression of translation, transcription, and cell-cycle arrest (**Hypothesis 3**). Furthermore, we expect: activation of anti-apoptotic pathways that keep cells alive at cold temperatures (**Hypothesis 4**), a systemic fuel switch from carbohydrates to lipids (**Hypothesis 5**), global suppression of metabolic genes in the ETC, OXPHOS, TCA cycle (**Hypothesis 6**), and altered or temperature-dependent RNA processing (**Hypothesis 7**). Ultimately, by identifying these pathways, we aim to determine whether hummingbirds utilize molecular pathways shared with other heterotherms, or if they have evolved a unique set of molecular strategies to achieve such extreme physiological plasticity. This approach could provide both mechanistic and evolutionary insights into how torpor is regulated in birds.

**Figure 1.**
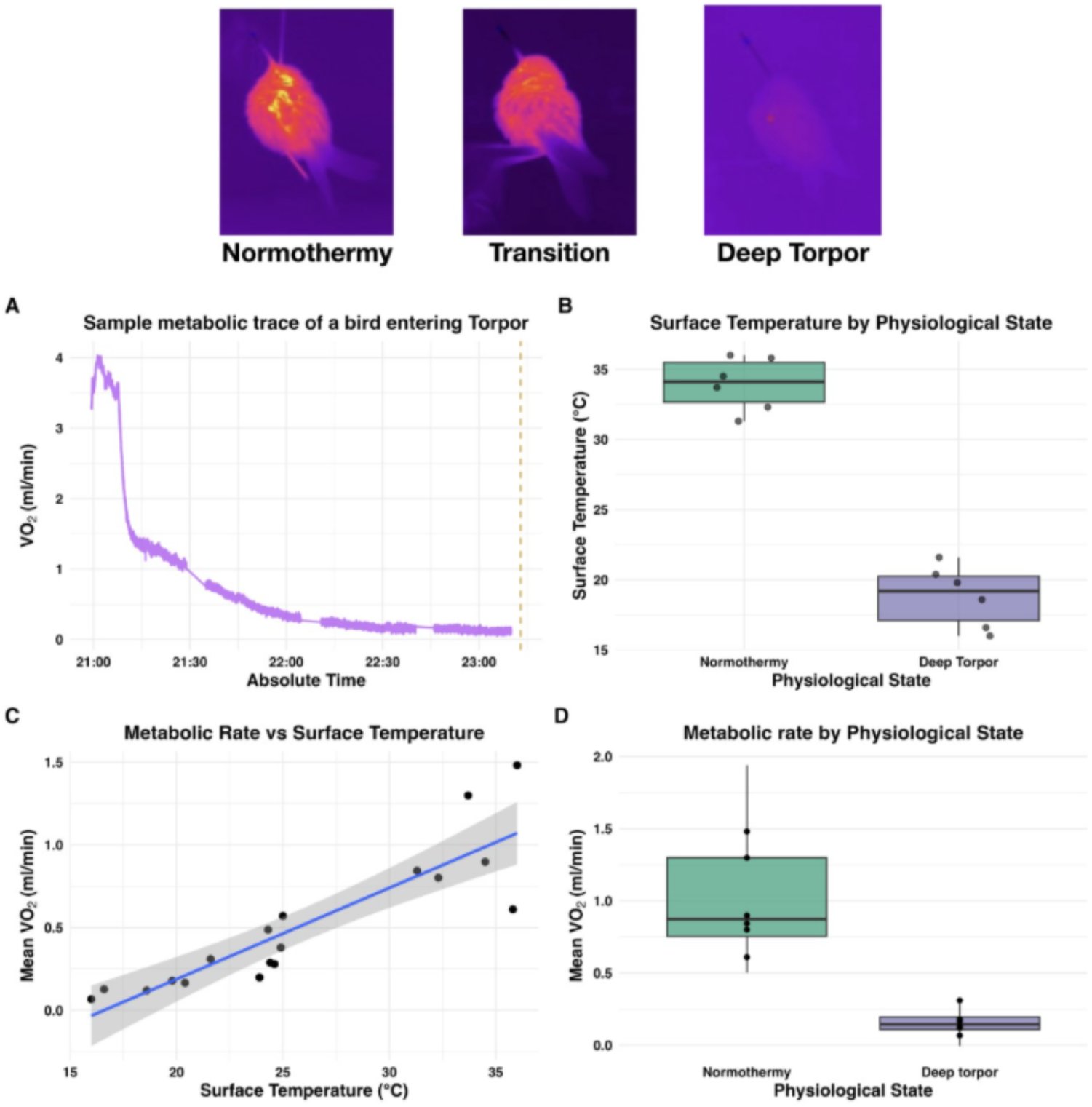
Modulation of whole organism metabolic rate and surface temperature in Anna’s hummingbirds (*Calypte anna*) during torpor entry. Top panel: Representative thermal images of birds in different physiological states adapted from (67). A: Representative metabolic rate trace of a bird entering torpor (gaps are baselining periods); B: Surface temperature of birds averaged over the last ten minutes before euthanasia with isoflurane; C: Correlation between metabolic rate and surface temperature of these birds (adjusted R² = 0.79, p< 0.001); D: Metabolic rates of birds averaged over the last ten minutes before euthanasia with isoflurane.

## Results

### Metabolic rate and surface temperature of birds in torpor

Surface temperature (t = 12.785, df = 9.77, p-value = 2.02e-07; Figure 1B) and metabolic rates (t = 5.944, df = 5.62, p-value = 1.27e-03; Figure 1D) were different across physiological states of normothermy and deep torpor. Birds took between 1 to 2.5hr to complete entry into torpor with body temperatures reaching within 3 degrees from the ambient (Figure 1A, Figure S1A, S1B). Mean VO₂ in the final ten minutes before euthanasia was highest in normothermy, intermediate in the non-stable state transition, lowest in deep torpor, and was significantly positively correlated with surface temperature (adjusted R² = 0.79, p< 0.001; Figure 1C). Birds in transition (classified using body temperature) exhibited substantial inter-individual variability in metabolic rate at similar surface temperatures (∼24–25 °C).

### Modelling expression of temperature-responsive genes

Transcriptomes of birds in transition resembled those of torpid birds more than those of normothermic birds (Figure 2A). To identify genes whose expression tracked physiological state or temperature, we screened 15,535 genes using a likelihood ratio test comparing a model with both state and temperature against an intercept only model (tissues pooled, bird ID as a random intercept; see Methods). This identified 151 responsive genes (p-adj < 0.05). This screen is different and pooled-across-tissues when compared to the per-tissue differential-expression contrasts below, and it has no fold-change thresholds. We classified the trajectories of these 151 genes by comparing four models using AIC. Thirty-seven genes (24.5%) were *linear*, tracking surface temperature continuously; 76 genes (50.3%) were *non-linear* meaning that state was a better descriptor of gene expression patterns. Of these 76, 37 were *early step genes* (changing between normothermy and transition), 12 were *late step genes* (changing between transition and torpor); and 27 were *graded/non-monotonic*, a pattern that neither a line nor a single step captured (Figure 2A). The remaining 38 genes (25.2%) were *indistinguishable* between a linear and a non-linear model (Figure 2C). To assess these proportions, we compared them with a permutation null. In each permutation, temperature was shuffled among birds within each state, which preserves state-level differences but removes any real temperature effects beyond state. The whole pipeline including the screen was repeated iteratively across the permutation runs. The *linear* fraction was higher than in the null (24.5% vs 14.5%, p = 0.01; Figure 2C), and so was the fraction of genes with a temperature effect beyond state (16.6% vs 8.8%, p = 0.01; Figure 2B), indicating a continuous temperature-dependent component. The *non-linear* fraction, however, was not higher than expected by chance (50.3% vs 45.2%, p = 0.14), so we do not interpret these genes as showing evidence of a discrete switch (Figure 2C). *Linear* genes included the splicing regulators SRSF1, SRSF2 and CLK4, whereas NR1D1, BHLHE40 and SGK1 were *graded/non-monotonic*. Repeating the analysis with log_10_ VO₂, instead of temperature, as the predictor gave the same pattern of excess within-state and *linear* signal over the null (Figure S18).

**Figure 2.**
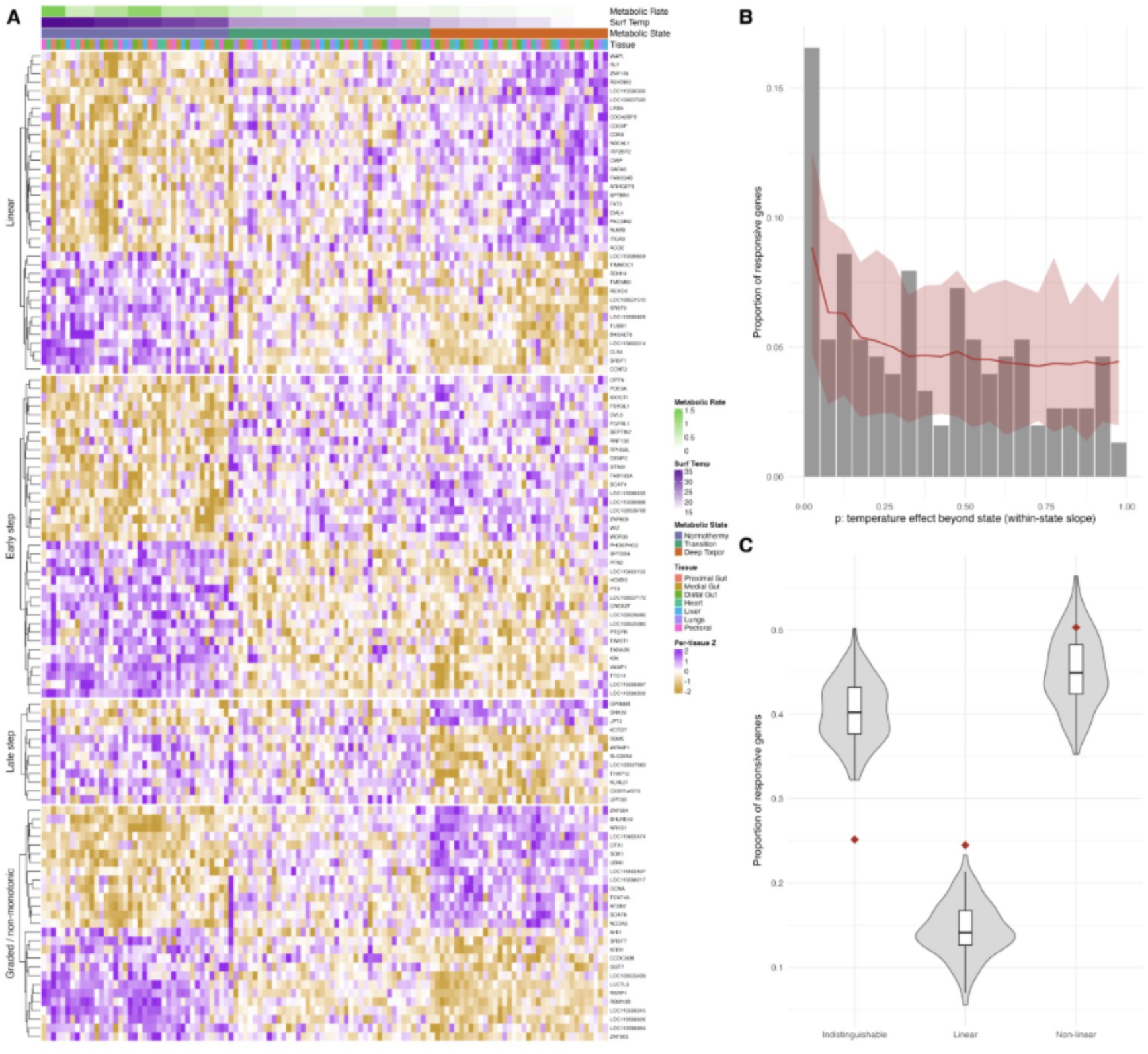
Most temperature-responsive genes differ between states; a smaller set tracks surface temperature continuously. **(A)** Expression of the 113 responsive genes classified as Linear (N = 37), Early step (N = 37), Late step (N = 12) or graded/non-monotonic (N = 27), grouped by class and clustered within class. Each column is one sample (bird × tissue), ordered from warmest to coolest surface temperature. Colours show per-tissue Z-scores of rlog-normalised expression. Top bars show metabolic rate, surface temperature, metabolic state and tissue. Indistinguishable genes (N = 38) are not shown. **(B)** Distribution of p-values for the temperature effect beyond state (state-means model vs state-means plus temperature) across the 151 responsive genes. Grey bars show the observed proportion of genes per bin (bin width 0.05). The red line and band show the mean and 95% range of 100 permutations in which surface temperature was shuffled among birds within state and the full analysis, including gene screening, was repeated. An excess of small p-values over the null indicates a continuous temperature component. **(C)** Proportion of responsive genes in each broad class (Linear, Non-linear, Indistinguishable). Violins and boxes show the permutation null (box: median and interquartile range). Red diamonds show the observed proportions. Linear genes exceed the null (24.5% vs 14.5%, p = 0.01), whereas non-linear genes do not (50.3% vs 45.2%, p = 0.14). Indistinguishable genes were fewer than in the null (25.2% vs 40.4%). Empirical p = (1 + number of permutations ≥ observed) / (1 + 100).

### Differential gene expression analyses

The degree of transcriptome remodelling in torpor varied across tissue types. Under strict differential expression thresholds (p-adjusted < 0.05 & log2FC > 0.58), the largest transcriptomic shifts were seen in the foregut, which saw roughly 4.4% (689 differentially expressed genes, or DEGs), and the liver, which saw 2.1% (320 DEGs) of the total genome differentially expressed. The heart, lung, and pectoral muscle showed comparatively little differential gene expression, which accounted for <1% of the genes tested (31 and 13 DEGs; Figures 3A, 3B). In the proximal and medial gut, downregulated genes vastly outnumbered upregulated ones, yielding a strong left skew in volcano plots with downregulated genes exhibiting large fold changes (on the order of log2 fold change ∼20; Figure 3A). Conversely, the liver displayed a right-skewed (more upregulated) volcano plot. The lungs, heart and pectoral muscles had very few DEGs, and their volcano plots were not skewed in either direction (Figure 3A). Although the majority of DEGs were tissue-specific with large sample-to-sample variance in expression (Figure S3), more than twenty genes were consistently differentially expressed between deep torpor and normothermic groups across three or more tissues. Seven of these genes (“Top Genes”) were shared DEGs across five or more tissue types and spanned distinct functional categories. These included transcriptional regulation and RNA splicing (RBM12B, RSRP1, and CLK4), as well as metabolic, circadian, and anti-apoptotic signalling (NR1D1 and SGK1). The remaining two genes were unannotated (LOC103537172 and LOC115598345; Figure 3B). Differential expression heatmaps revealed a conserved directionality of expression for these Top Genes across all organs except the lung (Figure 3B).

**Figure 3:**
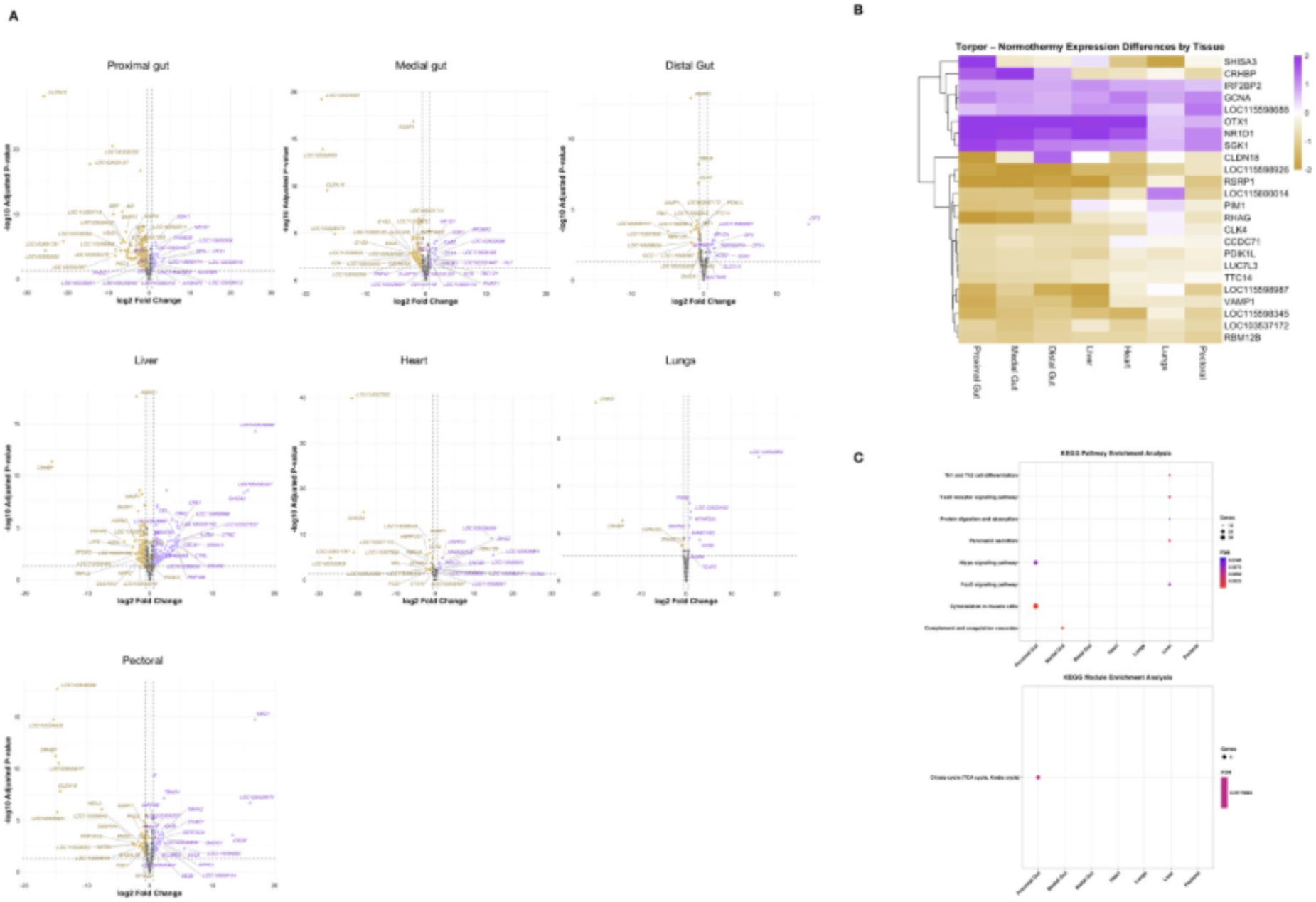
Differential expression analysis of Deep Torpor vs Normothermy indicates limited transcriptomic remodelling across tissue types. **(A)** Volcano plots showing genes upregulated in torpor (purple) and downregulated in torpor (yellow; p.adj < 0.05 & log2FC > 0.58). **(B)** Heatmaps showing ‘Top Genes’ differentially expressed in 5 or more tissue types. **(C)** KEGG pathway and module enrichments performed on the DEGs identified across tissue types.

Given the highly restricted number of strict DEGs, especially in all tissues other than the foregut and liver, we relaxed the framework by ignoring fold changes in expression (p-adjusted < 0.05 only) before running KEGG enrichments. These analyses revealed tissue-specific physiology with the proximal gut exhibiting cytoskeletal remodelling and TCA cycle modulation (Figure 3C). The liver displayed significant enrichment in FOXO signalling and immune-related pathways like T-cell receptor signalling (Figure 3C).

### Weighted Gene Correlation Network Analysis (WGCNA)

To move beyond individual gene thresholds and examine larger regulatory architecture, we used Weighted Gene Correlation Network Analysis (WGCNA). The complexity of transcriptomic change varied across tissues, with the highest number of distinct co-expression modules in the heart and lungs (Figure 4B, Figure S4). Module number reflects co-expression structure rather than the magnitude of change in torpor, so the modules associated with physiological state may capture small, coordinated shifts that individual-gene thresholds miss. KEGG enrichment of significant modules revealed tissue-specific regulation of processes. While the proximal gut was enriched in energy metabolism and lipid precursors via the TCA cycle and steroid biosynthesis, the medial gut and heart both exhibited significant enrichment in cytoskeletal remodelling, with the latter also involving AMPK signalling for energy homeostasis (Figure 4B). Notably, the pectoral muscle demonstrated a robust metabolic shift toward aerobic efficiency through enriched oxidative phosphorylation and TCA cycle pathways (Figure 4B, 4C), while the liver and lungs displayed more specialised responses in circadian rhythm regulation and proteostatic stress management, respectively (Figure 4B).

**Figure 4.**
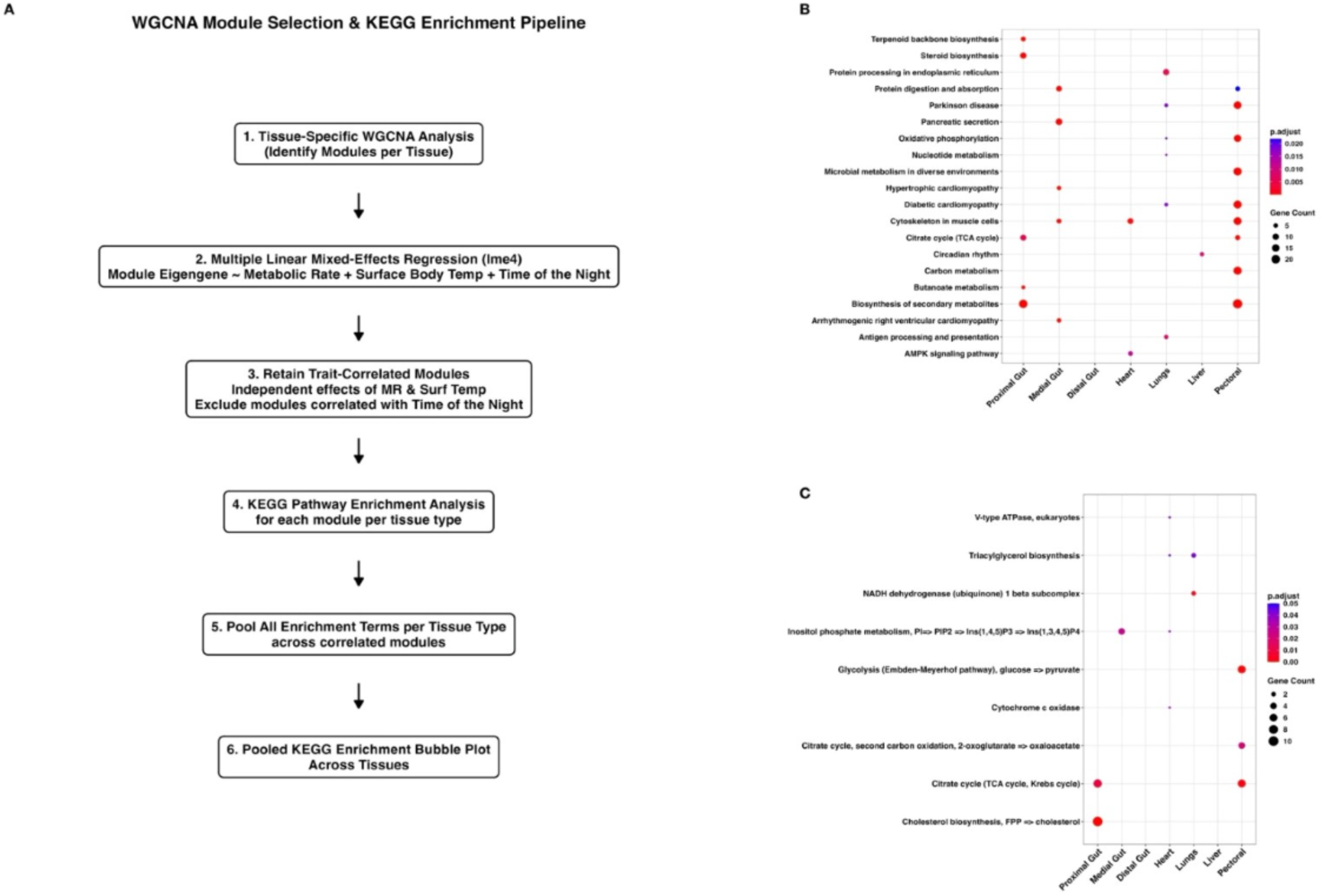
Tissue-specific transcriptional signatures of gene co-expression modules (WGCNA eigengenes) and physiological predictors across seven tissues. A: Flowchart describing the schematic followed to perform enrichments on modules relevant to avian torpor entry. B: KEGG pathways that were enriched across tissue types. C: KEGG modules that were enriched across tissue types.

### Targeted analysis of core molecular pathways

Examination of specific target pathways revealed differential expression patterns across tissue types (Table 1). Each molecular pathway that accounts for a large fraction of the cell’s energy was examined closely: about 28-34% of the cell’s ATP production goes to protein synthesis (estimated from mammalian cells and stimulated thymocytes), followed by about 25% to RNA and DNA synthesis - which are the most sensitive to decreases in energy supply (29, 31). Translation accounts for most of the cost of gene expression (30). Beyond translation and transcription, we examined processes previously hypothesized to be important in regulating mammal torpor: cell cycle and FOXO/Akt signalling, metabolism and thermogenesis, circadian regulation, and the spliceosome. We report the results in this order.

**Table 1.** Selected molecular players involved in early torpor of *Calypte anna*. This is a curated list meant to draw attention to key genes, not a complete list of changes; many more individual genes were significant when pooled across tissues and are reported in the Results and Figures S5–S15. The table shows the regulation of upstream and downstream genes, their known functions and how they affect organismal physiology and processes in different functional categories. Genes were selected in two ways. First, we tested whole pathways and curated gene sets (self-contained and competitive tests) and then examined which individual genes drove the sets that shifted. Second, we examined genes of known importance in torpor and basic cell biology individually, even where their set did not shift as a whole. Arrows show the direction of change in torpor relative to normothermy, pooled across the six tested tissues (lungs excluded) with bird identity as a random effect. Many of the listed genes are significant when pooled across tissues (p-adj < 0.05, Benjamini–Hochberg across the genes tested in each pathway); exact pooled p-adj values are given in the Results. “Trend” marks a consistent direction that did not reach p-adj < 0.05, either pooled across tissues (typically p-adj 0.05–0.15) or, where a tissue is named, within that tissue only. Tissue names in brackets indicate that the change was confined to that tissue.

| Functional Category | Upstream Genes & Functions | Downstream Genes & Functions | Likely effects |
|---|---|---|---|
| Global Transcription | GTF2E1↑, GTF2F2↑, GTF2H3↓ (GTF2E2↓ trend): general transcription factors.<br>MED6↓, MED4↓: Mediator subunits.<br>TAF3↑: TFIID subunit. | POLR2D↓ (RNA polymerase II), POLR3H↓ (RNA polymerase III), POLR1C↓ (RNA polymerase I): core polymerase subunits. | No coordinated change in the transcription machinery. A few subunits are lower and others higher, and the polymerase sets tend lower but not significantly. There is no sign of a global shutdown of the transcription machinery at the mRNA level. |
| Global Translation | EIF5↑: translation initiation factor.<br>EIF2AK2↓ (trend), EIF2AK3↑ (trend, gut): eIF2α stress kinases. | MRPS2↑, MRPS18A↑, MRPL22↑ (MRPS5↑, MRPS28↑ trend): mitochondrial ribosome transcripts.<br>RPL7L1↓: cytosolic ribosomal protein. | The translation machinery as a whole is unchanged at the mRNA level, with no evidence of a global cytosolic slowdown. Mitochondrial ribosome transcripts are higher across tissues, while cytosolic ribosome transcripts show only a weak, non-significant downward trend. |
| Cell Cycle & Anti-Apoptotic Signalling | SGK1↑, AKT1↑, PIK3R1↑: kinases and PI3K regulatory subunit, higher in all six tissues.<br>PDPK1↑ (trend), GSK3β↑ (trend).<br>AKT3: unchanged (paralog-specific response).<br>LAMC1↑, ITGA3↑: laminin chain and its integrin partner.<br>TLR4↑ (trend). | FOXO3↑: FoxO factor linked to cell survival.<br>FOXO1↓ (trend).<br>Cell cycle and repair gene set lower in all six tissues.<br>CDKN1A↓ (trend).<br>BCL2, CASP9: unchanged (apoptosis regulators). | Cells appear to shift toward protection and maintenance instead of growth. Survival signals (SGK1, AKT1) and a stress-protection factor (FOXO3) are higher, and genes for cell division and repair are slightly lower. Genes that control cell death did not change. |
| Metabolism & Thermogenesis | PDK3↓: PDH-inhibitory kinase; lower transcript suggests less restraint on PDH.<br>PRKACB↑, RPTOR↑: internal signalling anchors.<br>ADCY6↑ (trend).<br>RXRA↑ (trend, gut): upstream PPAR regulator. | DLAT↑, DLD↑: PDH subunits (entry to the cycle).<br>ACO2↑, IDH3B↑, SUCLG1↑, SDHA↑, SDHB↑, MDH2↑: core TCA enzymes.<br>NDUFS1↑, NDUFS2↑, NDUFV2↑ (complex I)<br>SDHA↑, SDHB↑ (complex II)<br>ATP5PF↑ (complex V)<br>SLC25A4↑ (ADP/ATP carrier), SLC25A3↑ (phosphate carrier)<br>MFN1↑, DNM1L↑ (trend): mitochondrial fusion and fission.<br>CPT2↑, ACADL↑, ACAA1↑, EHHADH↑ (trend, gut; not in liver): fatty acid oxidation.<br>PLN↓ (not significant): thermogenesis regulation. | Broad, modest upregulation of PDH, TCA and OXPHOS (complexes I, II and V) transcripts with lower PDK3, consistent with reduced restraint on PDH and increased TCA capacity. Mitochondrial dynamics and quality control set higher. No significant change in PPAR signalling or in fatty acid oxidation genes at the pathway level. |
| Circadian Regulation | CRY1↑, ARNTL↑, PER2↓, PER3↓: Core clock components regulating rhythm<br>GSK3β ↑: Kinase that phosphorylates and stabilizes Rev-erbα. | NR1D1 (Rev-erbα)↑, BHLHE40↑: Master transcriptional repressors.<br>G6PC↓ (liver): gluconeogenesis. NAMPT↑ (liver): NAD<sup>+</sup> salvage, clock-controlled. | Rewiring of the core clock with an upregulation of the repressor arm (NR1D1, BHLHE40, CRY1) and lower PER2/PER3. In the liver, lower G6PC is consistent with reduced glucose production. |
| Spliceosome Machinery | CLK1↓, CLK3↓, CLK4↓, RSRP1↓: SR-protein kinases and co-regulator.<br>HNRNPM↑: RNA-binding protein that can antagonise SR proteins. | SRSF1↓, SRSF2↓, SRSF3↓, SRSF7↓: SR protein family splicing regulators.<br>U2AF1↑, U2SURP↑, SF3B1↑, WBP11↑, PRPF8↑, DHX15↑: Branch-point recognition and core catalytic components. | Selective suppression of SR-protein splicing regulators and their kinases, with higher transcripts of core branchpoint and catalytic factors, while the spliceosome is unchanged. |

#### **a)** Global translation and transcription

Torpor did not globally change the expression of translation machinery (KEGG set, 177 KEGG ribosome and tRNA-synthetase genes present in our annotation: camera p-adj = 0.92; self-contained +0.04 Z, pooled lme p-adj = 0.80; Figure S5B), but two target sets shifted. First, the mitochondrial translation set was higher in all six tested tissues (+0.81 Z; camera p-adj = 0.29, not significant; self-contained, pooled lme p-adj = 0.013; mroast p-adj = 0.012), but was not significant in any single tissue (Figure S5B, 5C). This was driven by the mitochondrial ribosome transcripts MRPS2, MRPS18A and MRPL22 (pooled p-adj = 0.014, 0.020 and 0.020, each higher in all six tissues; Figure 5A), with MRPS5 and MRPS28 tending higher (p-adj = 0.084 and 0.092). Second, the initiation factor set was also higher (+0.39 Z; camera p-adj = 0.77, not significant; self-contained p-adj = 0.031; Figure S5B), driven by EIF5 (p-adj = 0.020; Figure S5A), which was highest in the gut segments (Table 1). The selected cytosolic ribosomal genes tended lower (−0.57 Z; camera p-adj = 0.77; self-contained p-adj = 0.14, a weak trend), driven by RPL7L1 (p-adj = 0.008, lower in all six tissues), whereas RPL29 and RPL34 did not change (Figure S5A). The stress-kinase set was flat overall (camera p-adj = 0.92; self-contained p-adj = 0.65) but its members moved in opposite directions (mroast p-adj = 0.03; Figure S5B), with EIF2AK2 tending lower (p-adj = 0.084**; p-nominal = 0.021**) and EIF2AK3 higher in the proximal and medial gut only before correction (p-nominal = 0.014 and 0.0025; p-adj = 0.30 and 0.16; Figure S5A).

**Figure 5.**
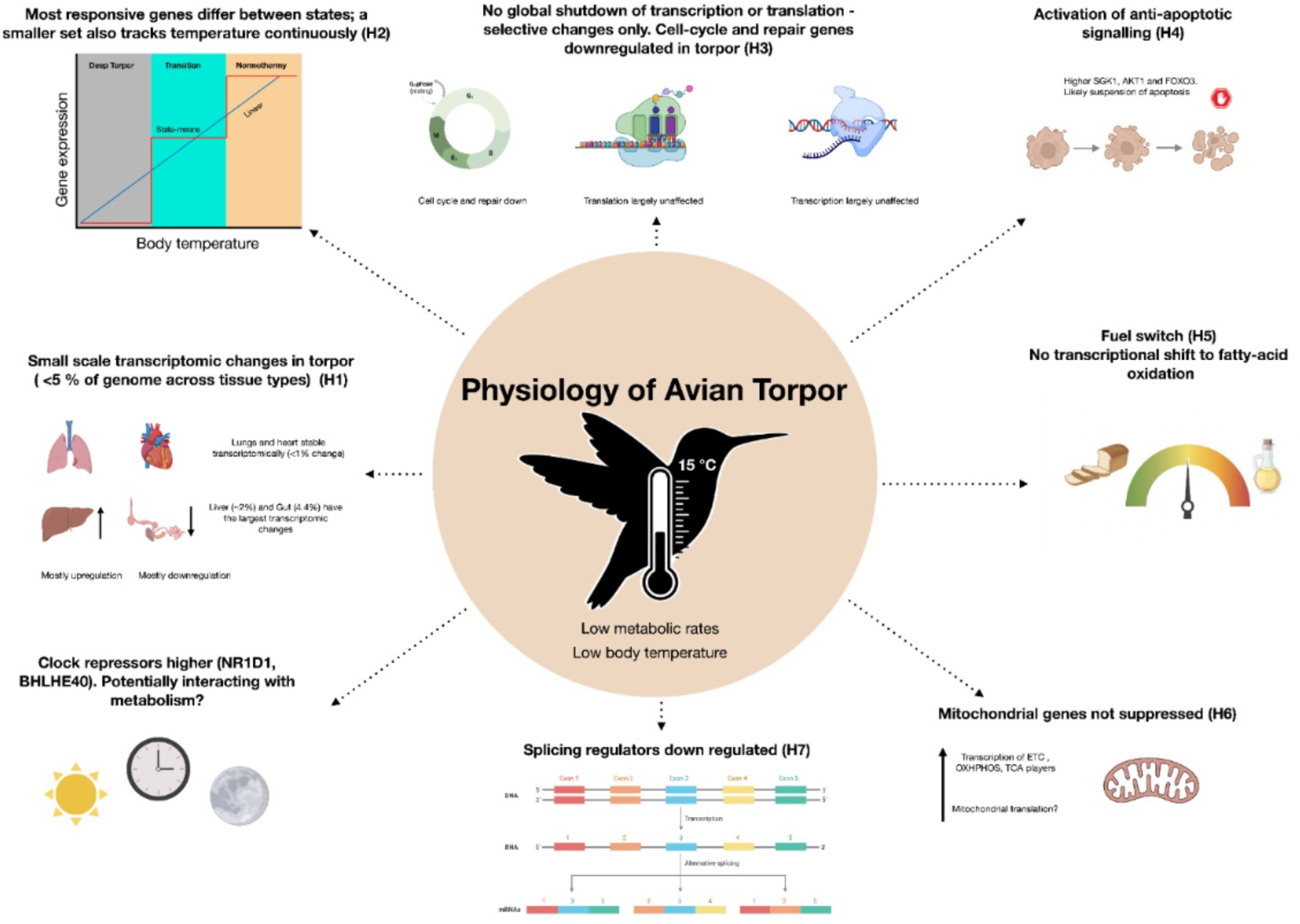
Summary of the paper with evidence for hypotheses outlined in the study. Some of the art was created in BioRender.

The expression of the transcription machinery did not change significantly. The whole KEGG set (51 of 78 genes present) and the three RNA polymerase sets all tended lower in torpor, with the whole set lower in all six tested tissues (−0.14 Z) and RNA polymerase III the most consistent (−0.29 Z, lower in 6/6), but none was significant in either test (camera p-adj ≥ 0.24; self-contained, pooled lme p-adj = 0.36; Figure S6B, 6C); mroast was significant for the whole set (p-adj = 0.0035) and for RNA polymerase III (p-adj = 0.0056). The polymerase signal came from single subunits: POLR2D (RNA polymerase II), POLR3H (III) and POLR1C (I) were lower in all six tissues (pooled p-adj = 0.0004, 0.0007 and 0.0495; Figure S6A), although none was significant in any single tissue after correction. The general transcription factor, Mediator and TFIID sets were also unchanged on average (camera p-adj ≥ 0.63; self-contained p-adj ≥ 0.43), but their members moved in both directions (mroast p-adj = 0.0035 for each; Figure S6B). Among the general factors, GTF2E1 and GTF2F2 were higher when expression profiles were pooled across tissue types (p-adj = 0.022 each) and GTF2H3 was lower (p-adj = 0.022), with GTF2E2, the partner of GTF2E1, tending lower (p-adj = 0.053**; p-nominal = 0.0088**; Figure S6A). In Mediator, MED6 and MED4 were lower in all tissues (p-adj = 0.0066 and 0.022), whereas MED14 and MED17 were higher only in parts of the gut and not after correction (pooled p-adj = 0.14 and 0.20**; pooled p-nominal = 0.032 and 0.066**; Figure S6A). Among TFIID subunits, TAF3 was higher (p-adj = 0.011; Figure S6A).

#### **b)** Cell cycle, PI3K-AKT, and FOXO signalling

Torpor was accompanied by higher expression of the Akt/SGK kinase module and a modest decrease of cell-cycle and repair genes (Figure S8) without a major reorganization of cell cycle players (Figure S7). The kinase module shifted up (camera p-adj = 0.16; self-contained, pooled p-adj = 0.069, trend; mroast p-adj = 0.005; Figure S8B). SGK1 drove this (pooled p-adj < 10⁻⁷; significant after correction in proximal gut, medial gut, heart and pectoral; Figure S8A). AKT1 (p-adj = 0.011) and the PI3K regulatory subunit PIK3R1 (p-adj = 0.006) were also upregulated in all six tissues, PDPK1 followed as a trend (p-adj = 0.098**; p-nominal = 0.0095**), and AKT3 was unchanged (p-adj = 0.47), so the response is paralog-specific (Figure S8A). Within the ECM and integrins set, which was not significant as a whole (camera p-adj = 0.71; self-contained p-adj = 0.055; Figure S8B, 8C), the laminin chain LAMC1 and its integrin partner ITGA3 were each higher in all six tissues (both pooled p-adj = 0.002; Figure S8A).

The FoxO factors moved in opposite directions: FOXO3 rose in torpor in all six tissues (p-adj = 2×10⁻⁴), while FOXO1 tended to decrease (p-adj = 0.10–0.13**; p-nominal = 0.014**, strongest in heart and liver; Figure S9A). This is consistent with FOXO3’s canonical role in suppressing apoptosis and promoting cell viability (32). After accounting for bird identity as a random effect, AKT1 expression was not significantly associated with FOXO3 (bird-aware p-nominal = 0.059) or FOXO1 (p-nominal = 0.114), and AKT3 was not associated with SGK1 (p-nominal = 0.76). GSK3B rose in all six tissues as a trend (p-adj = 0.08**; p-nominal = 0.0070**; Figure S8A). Receptor-level changes were weaker: TLR4 was up in all six tissues as a trend (p-adj = 0.079**; p-nominal = 0.0050**) and PTK2 only nominally (p-adj = 0.20; p-nominal = 0.026; Figure S8A). Cell-cycle genes drifted down: the cell cycle and repair set was lower in all six tissues (camera p-adj = 0.16; self-contained, pooled p-adj = 0.037**; mroast p-adj = 0.020**, largest in proximal gut; Figure S8B). No single member passed the threshold, and CDKN1A (p-adj = 0.098**; p-nominal = 0.0099**) came closest (Figure S8A). Apoptosis regulators were not differentially expressed (BCL2 p-adj = 0.78, CASP9 p-adj = 0.63; Figure S8A).

#### **c)** Metabolism (TCA, OXPHOS, ETC, PPAR) and thermogenesis

Although torpor is a state of metabolic suppression, TCA cycle transcripts were modestly but consistently higher in torpid birds. Competitive (camera) tests were not significant after correction (pooled p-adj = 0.076 for the whole KEGG TCA pathway and 0.076–0.129 across its units; Figure S10C). Set-level self-contained tests showed that the whole KEGG TCA pathway shifted upward in torpor when pooled across tissues (+0.65 Z; pooled lme, BH p-adj < 0.01; Figure S10B), supported by the any-direction (mroast) test (p-adj = 0.009; 0.006–0.013 across the units). The shift was strongest in the liver and gut, but no tissue reached significance on its own (Table 1). The shift was carried by all four KEGG functional units (Figure S10B), whereas the non-KEGG pyruvate entry and PDH regulation set did not shift (−0.09 Z; self-contained p-adj = 0.47; Figure S10B). The citrate and isocitrate steps (+0.73 Z, p-adj < 0.01) were led by ACO2 (p-adj < 0.01) and IDH3B (p-adj < 0.05), and the α-ketoglutarate and succinate node (+0.54 Z, p-adj < 0.01) by SUCLG1 (p-adj < 0.01). In the succinate-to-oxaloacetate segment (+0.76 Z, p-adj < 0.01), SDHA, SDHB and MDH2 were higher (p-adj < 0.05). The pyruvate dehydrogenase (PDH) complex (+0.71 Z, p-adj < 0.05) showed higher DLAT and DLD (p-adj < 0.05) and lower PDH kinase PDK3 (pooled p-adj = 0.028; lower in five of six tissues, with distal gut and liver p-nominal = 0.042 and 0.023; Figure S10A). All these genes were higher in at least five of six tissues, and none was significant within a single tissue (BH across the 31 TCA genes; Figure S10A). Together, these changes are consistent with reduced inhibition of PDH and increased TCA capacity.

OXPHOS transcripts showed a similar modest, broad upward shift. Competitive (camera) tests were not significant (p-adj = 0.23 for the whole pathway and for each of complexes I, II and V; Figure S11C). Pooled across tissues, the whole KEGG OXPHOS pathway shifted upward (+0.48 Z; pooled lme, BH p-adj < 0.01; Figure S11B), supported by the self-contained test of each complex and the any-direction (mroast) test (p-adj = 0.009). The shift was in the same direction in every tissue, strongest in the liver and distal gut, although no tissue reached significance alone. Complexes III and IV, and mitochondrially encoded subunits, could not be assessed because these genes are absent from the processed expression matrix; the analyses therefore cover complexes I, II and V (the whole-pathway test includes 72 of 138 KEGG OXPHOS genes). Complex I was enriched (+0.62 Z, p-adj < 0.01**; mroast p-adj = 0.024**; Figure S11B) and supported by the upregulation of NDUFS1, NDUFS2, NDUFV2, NDUFA9, NDUFA10 and NDUFV3, and complex II (+0.88 Z, p-adj < 0.01**; mroast p-adj = 0.009**; Figure S11B) by SDHA and SDHB (pooled p-adj < 0.05, BH across the 72 OXPHOS genes; Figure S11A). Complex V (+0.61 Z, p-adj < 0.05**; mroast p-adj = 0.034**) showed a consistent upward shift across all 13 of its genes, with ATP5PF individually significant (p-adj = 0.040) (Figure S11A).

PPAR signalling, which has major roles in fatty acid oxidation, lipid transport and crosstalk with insulin signalling (33, 34), showed no significant pathway-level change in torpor, either in any individual tissue or when pooled across tissues (camera p-adj ≥ 0.56; self-contained not significant; mroast p-adj ≥ 0.19; Figure S12B, 12C). Among the fatty-acid oxidation genes, a modest trend towards higher expression (RXRA, CPT2, ACADL, ACAA1, EHHADH, ACOX1) was visible in the gut but did not reach significance, whereas in the liver these genes did not change or were lower (CPT1B−1.2 Z, p-adj = 0.62; p-nominal = 0.027; CPT1A, EHHADH and ACAA1 −0.3 to −0.5 Z; Figure S12A). Among the lipoprotein-related genes, only PLTP was higher in torpor in every tested tissue (pooled p-adj = 0.41; p-nominal = 0.017). In the liver, ADIPOQ transcripts were higher (+1.4 Z; p-nominal = 0.0045) but did not survive correction (BH p-adj = 0.62), and the liver fatty-acid oxidation genes above did not follow (Figure S12A) (33).

Of the ten thermogenesis-related gene sets, none shifted significantly in any single tissue. Pooled across tissues, the mitochondrial dynamics and quality control set was higher in torpor in all six tested tissues (+0.39 Z; camera p-adj = 0.23; self-contained BH p-adj = 0.043**; mroast p-adj = 0.043**; Figure S13B), with the fusion and fission genes MFN1 and DNM1L trending higher (p-adj = 0.08 and 0.07; p-nominal = 0.0053 and 0.0040; Figure S13A) (34). The whole KEGG thermogenesis pathway was also higher (+0.30 Z; camera p-adj = 0.23; self-contained p-adj = 0.043**; mroast p-adj = 0.025**), but this largely reflects its OXPHOS genes: without them it was not significant in the self-contained test (p-adj = 0.31**; camera p-adj = 0.23), although mroast remained significant (p-adj = 0.010); Figure S13B, 13C). The adrenergic and mTOR signalling sets did not shift significantly, although PRKACB and RPTOR were individually higher when pooled (p-adj = 0.036 and 0.014; Figure S13A). A similar trend was observed with ADCY6 (p-adj = 0.07; p-nominal = 0.0029**). In the calcium-cycling and uncoupling set, PLN was lower in all six tested tissues but not significantly (p-adj = 0.27**; p-nominal = 0.022**), which at most hints at a slight withdrawal of SERCA-associated non-shivering thermogenesis (35) (Figure S13A). In the antioxidant set, TXNRD1 was lower in the liver (p-adj = 0.58; p-nominal = 0.033; Figure S13A), but not after correction (36), and, surprisingly, GPX4 and other negative regulators of ferroptosis were not significantly upregulated, unlike in hibernator cell lines under cold stress (15, 17).

#### **d)** Circadian regulation

We found that core clock genes were rewired in torpor despite controlling for time of the night at which the birds across conditions were sampled (Figure S14). The clock gene sets (core timekeepers, clock kinases and F-box proteins, and the whole KEGG circadian pathway) changed significantly in the any-direction mroast test (p-adj ≤ 0.006) but not in the directional self-contained test (p-adj ≥ 0.39; Figure S14B), because their members moved in opposite directions. Within the core timekeepers, ARNTL (BMAL1) and the repressor CRY1 were higher in all six tested tissues (pooled BH p-adj = 0.017 and 0.001), whereas PER2 and PER3 were lower in all six (p-adj = 0.004 and 0.0009; Figure S14A) (37). After per-tissue correction, CRY1 was significant in the pectoral muscle only (Figure S14A). The set of genes linking circadian signalling to metabolism shifted more than other genes when pooled (+0.41 Z; camera BH p-adj = 0.048), driven by the repressors of CLOCK–BMAL1 activity NR1D1 (Rev-erbα) and BHLHE40 (Figure S14A, 14C). Both genes were higher in all six tissues (pooled p-adj = 2×10⁻¹² and 4×10⁻⁷) in torpor and are known to actively suppress metabolic gene expression (38, 39). NR1D1 was significant in every tested tissue and BHLHE40 in the heart, liver and pectoral muscle. GSK3B, which stabilises Rev-erbα, was also higher in torpor (p-adj = 0.041; Figure S14A), as was the clock-controlled NAD⁺ salvage gene NAMPT in the liver (p-adj = 0.031; Figure S14A). The fuel-selection and energy and stress monitoring sets did not shift (camera p-adj ≥ 0.86; Figure S14C), and evidence for repression of metabolic targets was confined to the liver, where G6PC was lower (p-adj = 0.028), consistent with Rev-erbα repression of gluconeogenesis (33). However, PPARG, APOC3, PDK4 and PPARGC1A did not change significantly after correction (Figure S14A) (33).

#### **e)** Spliceosome

We found that torpor selectively silences the splicing regulatory machinery (Figure S15). The whole KEGG spliceosome pathway and the core snRNP/Sm proteins were unchanged (pooled lme p-adj = 0.99), but the set of SR proteins and splicing regulators were lower in all six tested tissues (−0.50 Z; pooled lme BH p-adj = 0.004; camera p-adj = 0.043; Figure S15B, 15C), although no single tissue reached significance. This was driven by the SR proteins SRSF1, SRSF2, SRSF3 and SRSF7 (pooled p-adj = 3×10⁻⁸, 4×10⁻⁷, 0.014 and 0.007), the SR-protein kinases CLK1, CLK3 and CLK4 (p-adj = 2×10⁻⁵, 0.007 and 10⁻¹⁴), and by their co-regulator RSRP1 (p-adj = 2×10⁻¹⁸), which was significantly lower in every tested tissue (Figure S15A). SRSF1 and SRSF7 were also individually significant in the medial and distal gut, CLK4 in all tissues except the pectoral muscle, and SRSF2 in the liver (Table 1). In the same set, the RNA-binding protein HNRNPM, which can antagonise SR proteins, was higher in all six tissues (+1.1 Z; p-adj = 0.002; Figure S15A)(40). In contrast, the sets for branch-point recognition, helicases and assembly, and other associated factors tended higher (+0.24 to +0.32 Z; p-adj = 0.075), driven by U2AF1, U2SURP and SF3B1(41), by WBP11, and by the catalytic core components PRPF8 and DHX15 (all higher in every tested tissue; p-adj = 0.0004–0.045)(42), whereas PRPF38B was lower (p-adj = 0.0008; Figure S15A).

## Discussion

We set out to examine the molecular mechanisms and tissue-specific regulatory patterns that allow hummingbirds to switch from a state of normothermy to early torpor. To the best of our knowledge, this represents the first whole-transcriptomic characterization of torpor in birds. By studying torpor across a range of endothermic vertebrates (birds vs. mammals), we can better understand the evolutionary history of endothermy and heterothermy. We found that 1–4.4% of the genes tested were differentially expressed across tissue types in hummingbirds. For comparison, the most responsive tissue reached 1.0% in the Djungarian hamster (hypothalamus, torpor entry), 1.7% in the grey mouse lemur (skeletal muscle, miRNA) and 6.7% in the mouse (soleus muscle, promoters) among species using daily or fasting-induced torpor. Among hibernators it reached 1.8% in the fat-tailed dwarf lemur (white adipose tissue), 11.3% in the greater horseshoe bat (wing), 17.2% in the arctic ground squirrel (liver) and 34.6% in the 13-lined ground squirrel (brown adipose tissue) (Table S1). However, these values are not strictly comparable because the number of genes tested, the statistical thresholds and the contrasts differ among studies. Despite this, the patterns of differential expression suggest perhaps that the degree of transcriptional change in avian daily torpor parallels mammalian daily torpor, as do the time scales, while they are lower than most mammalian hibernators. A set of seven genes seemed consistently differentially expressed across three or more tissue types, but the majority of the other DEGs were only different in one tissue type each. Energetically costly processes were not globally suppressed, suggesting regulated changes in RNA processing rather than a general shutdown of cellular machinery.

Our study supports some of our *a priori* hypotheses but not others. We found that, contrary to our expectations, there is very limited transcriptome remodelling in torpor (Hypothesis 1). We find that most temperature-responsive genes differ between physiological states, and a smaller set tracks temperature continuously more than expected by chance, but we find no evidence of a discrete switch (Hypothesis 2). We found no evidence of global transcriptional or translational suppression, only a modest drift down of cell cycle and repair genes (Hypothesis 3), and apoptosis regulators did not change at a large scale, although higher SGK1, AKT1 and FOXO3 are consistent with survival signalling (Hypothesis 4). We found no transcriptional signature of a shift toward lipid metabolism (Hypothesis 5) despite evidence for this at the organismal level. We also find that rather than a suppression of mitochondrial genes as expected, there is instead a modest upregulation of TCA cycle and OXPHOS genes at the transcript level (Hypothesis 6). We do document altered RNA splicing regulation (Hypothesis 7). Interestingly, core clock genes and metabolism modulators were also rewired.

We found very few differentially expressed genes (1–4.4% of the genes tested) across tissue types, despite birds taking ∼60–150 minutes to complete entry into torpor, and despite using low thresholds for DEG discovery (Log2FC ≥ 0.58, p.adj ≤ 0.05). This suggests that the transition into torpor is either largely orchestrated at another molecular level (e.g., protein modifications or degradation) or that it is accompanied by targeted changes in a relatively small subset of transcripts that have large consequences for metabolism. This restricted remodelling of the transcriptome plays out differently across different tissue types. The proximal and medial gut show a negative skew in gene expression (i.e., more downregulation when compared to upregulation), while the liver exhibits a positive skew, and other tissues examined appear to be stable (i.e., approximately as many genes are downregulated as upregulated; as seen in the volcano plots, Figure 3). The gut, an energy-demanding tissue, had the largest set of downregulated genes. Furthermore, the degree of downregulation appears to be far greater when compared to the degree of upregulation observed across tissue types. Given the time scales of torpor entry (Figure S1) and previously published studies on hibernation, it is likely that strongly downregulated genes may reflect preferential degradation, whereas transcripts that remain abundant may be those that are selectively stabilised (23, 43). While RBM3 is implicated in stabilizing transcripts at low temperatures in ground squirrel hibernation (44), we found RBM3 was not present in our dataset at all, but that RBM12B was in many tissue types. RBM12B’s role in stabilising transcripts is poorly studied; direct measurements of RNA turnover would be needed to test this hypothesis.

Studies that have managed to measure organismal metabolic rates and body temperatures concurrently seem to suggest that torpor entry in mammals is initiated first with a drop in metabolic rate followed by a drop in body temperature (12, 45, 46). Our transition birds transcriptionally resembled torpid birds more than normothermic birds, and early step genes outnumbered late step genes (37 vs 12; Figure 2A). However, step-like genes were not more common than expected by chance (Figure 2C), so we do not interpret this as a discrete switch. Instead, we find a disproportionately higher proportion of genes that behaved linearly, more so than expected by chance. Part of the transcriptional response is therefore already under way by mid-transition, but our design cannot tell whether transcriptional change precedes or follows the metabolic rate changes.

While we recovered a handful of differentially expressed genes shared across tissue types (see ‘Top Genes’), most differentially expressed genes were tissue-specific. Furthermore, enrichment analyses identified very generic pathways that were not indicative of underlying physiology, probably due to limited transcriptional remodelling coupled with only a handful of key molecular players crossing the Log₂FC thresholds. Abandoning the Log₂FC thresholds and examining specific genes central to key pathways, while accounting for differences in basal levels of expression of genes across tissue types, reveals a selective, multi-tier molecular response. We discuss this response one pathway at a time with the sets that shifted as a whole (self-contained and competitive tests) first, the genes that drive them second, and finally the individual genes that shifted strongly although their sets did not.

The strongest pathway-level change was in splicing regulation: the set of SR proteins and splicing regulators was lower in all six tested tissues (−0.50 Z; pooled p-adj = 0.004 in the self-contained test and 0.043 in the competitive test; Figure S15). Hummingbirds in torpor exhibit a selective downregulation of key splicing regulators (SRSF1, SRSF2, SRSF3, SRSF7, CLK1, CLK3, CLK4 and RSRP1; Figure S15), seemingly like how hibernating ground squirrels and hamsters modulate their splicing landscapes by altering CIRP (cold-inducible RNA-binding protein) and SRSFs (47, 48). Suppressing these regulators may serve as a brake against aberrant splicing given that CLK4 activity increases at lower temperatures (49). Hibernating mammals accrue a sleep debt (12, 50, 51), and sleep deprivation increases RNA splicing activity in the mammalian hippocampus (52); if torpor imposes a similar sleep debt in hummingbirds, we might therefore have expected splicing regulators to rise, not fall (12, 50). Such changes in splicing regulators may result in altered splicing landscapes that have consequences for underlying physiology, and though we did not measure splicing directly in this study, these avenues are worth future exploration.

We noted an upregulation of mitochondrial ribosome subunits, namely MRPS2, MRPS18A and MRPL22, which was consistent across all six tested tissues (Figure S5). Furthermore, we found a counterintuitive modest upregulation of genes in the tricarboxylic acid cycle (DLAT, DLD, ACO2, IDH3B, SUCLG1, MDH2) and oxidative phosphorylation (NDUFs, SDHA, SDHB, ATP5PF) when pooled across tissues, although no single tissue was significant alone and complexes III and IV could not be assessed (Figures S10, S11; Hypothesis 6). We interpret this counterintuitive upregulation (i.e., mitochondrial transcriptional upregulation despite metabolic rate reduction) by drawing heavily on empirically tested models of hibernating mammals (7, 46, 53). We propose three distinct but potentially synergistic, hypotheses to explain this unexpected transcriptomic signature: a) Q10 effects—most torpid animals maintain body temperatures just a few degrees above ambient. Maintaining even low metabolic activity at these low body temperatures is incredibly slow and inefficient due to depressed enzyme kinetics. Under this model, the widespread upregulation of metabolic machinery might be a necessary compensatory mechanism to maintain activity at low body temperatures; b) Mitochondrial stability—the upregulation of Complex I and Complex II components may serve to keep mitochondria structurally stable at cold temperatures; c) Preparing for energetically demanding arousal. Many TCA cycle enzymes such as PDH are catalytically inhibited by phosphorylation (54). These enzymes are potentially paused in torpor, and metabolic precursors derived from slow breakdown of fat and amino acid breakdown stockpile in the mitochondria. Throughout the torpor bout, local ATP is slowly depleted for basal cellular maintenance, eventually triggering the dephosphorylation (reactivation) of these enzymes upon arousal. Our finding of lower PDK3, a PDH-inhibitory kinase, points the other direction (less restraint on PDH), so this third mechanism remains speculative. ADIPOQ transcripts were higher in the liver, but only nominally (p = 0.0045; p-adj = 0.62), and PPAR signalling and fatty acid oxidation genes showed no significant pathway-level change. We therefore have no transcriptional evidence for enhanced fatty acid oxidation in torpor, although lipid use during torpor has been shown at the organismal scale in hummingbirds and hibernating mammals (4, 33, 55, 56).

The circadian clock appears to be rewired in torpor, with prominent changes in transcriptional repressors (39). The circadian pathway changed in both directions, which is why it was detected by the any-direction test and not the self-contained test: the master timekeepers moved in opposing directions (CRY1 and ARNTL higher, PER2 and PER3 lower; Figure S14B, 14C), and the circadian–metabolic linkers were higher, particularly in the liver and proximal gut. This occurred despite the experiment actively controlling for the time of sampling of birds across metabolic states (see Methods). NR1D1 (Rev-erbα) and BHLHE40, which are major transcriptional repressors implicated in modulating metabolism (38, 57), were both higher in torpor; NR1D1 in all six tissues and BHLHE40 in the heart, liver and pectoral muscle (Figure S14A). In the liver, the only tissue where it changed, expression of G6PC, which encodes the enzyme that releases glucose from the liver, was lower. This is consistent with repression by NR1D1, a known repressor of hepatic gluconeogenic genes including G6PC (57), although we did not measure glucose output. Additionally, torpor is neurologically distinct from deep sleep (12, 50), and while both sleep deprivation and torpor severely alter molecular clocks (51), they elicit different metabolic responses: sleep deprivation involves upregulating metabolic genes to sustain high energetic demand (52, 58, 59), whereas in torpor metabolism is suppressed. A rewired clock with strong repressor signals fits this contrast and suggests that torpor may recruit the clock as a metabolic brake rather than simply disrupting it, although we did not measure the activity of these factors.

The differential expression of key targets involved in regulation of cell cycle and survival (higher SGK1, AKT1, PIK3R1 and FOXO3, unchanged AKT3, and consistently lower expression of the PI3K–Akt “cell cycle and repair” gene set, −0.36 Z pooled, p-adj < 0.05, lower in all six tissues, and of FoxO cell-cycle-arrest targets, whereas the whole KEGG cell cycle pathway did not change (Figures S7–S9). This paints a picture of a shift away from proliferation towards cell maintenance, not outright cell cycle arrest (54, 60, 61). Apoptosis regulators (BCL2, CASP9) and the apoptosis gene sets did not change, so these data neither support nor exclude a role for apoptosis in torpor: low temperature could itself promote or suppress apoptosis, and testing this would need direct measures of cell death. The paralog-specific response of AKT family members (AKT1 higher, AKT3 unchanged) has, to our knowledge, not been described in torpor, although Akt signalling is regulated in a tissue-specific way in torpid speckled mousebirds (8). We hypothesise that higher Akt/SGK signalling supports cell survival and tissue maintenance through two possible mechanisms: a) a stronger cell–matrix connection (LAMC1 and ITGA3 higher in all six tissues; Figure S8A) and b) SGK1 upregulation, which is known to prevent atrophy of skeletal muscles in hibernating ground squirrels (62–64).

Our strongest *a priori* prediction, that the more expensive cellular processes would be shut down, was not supported. Protein synthesis is the largest biosynthetic energy cost: about 28% of ATP production in mammals (29) and 34% of ATP consumption in stimulated thymocytes, where RNA and DNA synthesis use a further 25% (31). Per gene, translation accounts for about 94% of the cost of expression in *E. coli* and yeast, transcription for about 3–6%, and replication for 2% or less (30). We found no evidence of global suppression of these processes at the mRNA level. For translation, the core machinery was unchanged (apart from the mitochondrial translation set, above): the initiation factor set was higher (driven by EIF5), and cytosolic ribosomal genes showed only a weak, non-significant downward trend driven by RPL7L1, while RPL29 and RPL34 did not change (Figure S5). That cytosolic ribosomal transcription is suppressed is well-documented in mammalian hibernation, so our result differs from mammalian hibernation, although we cannot rule out suppression at the level of translation (3, 65). For transcription, global suppression is reported in hibernation in ground squirrels (3, 9), but we did not see it at the mRNA level. Individual RNA polymerase subunit genes (POLR2D, POLR3H and POLR1C) were lower in all six tissues, but the whole transcription machinery and the polymerase sets only tended lower and did not change significantly, while mediator and general transcription factors moved in both directions (Figure S6).

While this study provides novel insights into avian torpor through a molecular lens, several limitations in this study warrant close consideration. First, the physiological comparison in this study represents a circadian-controlled contrast between normothermy and early torpor. Sampling of birds when active (during the day) and at a later time point into torpor are likely to show a full range of change in gene expression patterns and more starkly highlight preparatory transcriptomic changes in hummingbirds prior to sleep. There is some evidence to suggest that synthesis of new transcripts in the Golden hamster, *Mesocricetus auratus*, is strongly dependent on the amount of time spent in different physiological states (66). Second, many of our functional interpretations are based on transcript abundance and may not directly reflect changes in protein abundance or activity. Furthermore, many proteins like CLKs and GSK3β operate by phosphorylating other proteins to activate their functions, and we have not assessed post-translational modifications. Third, transcript abundance estimates recovered following STAR alignment were not independently validated using qPCR. Fourth, lungs (fewer than three torpid birds) were excluded from the targeted tests, and OXPHOS complexes III and IV and mitochondrially-encoded genes were absent from the expression matrix. Finally, given the broad-scale nature of these analyses and our hypotheses, we warrant caution when interpreting individual genes and pathways. While numerous genes exhibited nominally significant differences in expression between physiological states within tissues, relatively few remained significant within a single tissue and most of our conclusions rest on effects pooled across tissue types. Consequently, many of the mechanistic interpretations proposed here should be viewed as hypotheses generated from coordinated patterns across genes, pathways, and tissues rather than definitive evidence for specific regulatory mechanisms. Future studies with complementary proteomic and phosphoproteomic datasets, and direct functional validation, will be necessary to further evaluate these hypotheses.

Despite the small degree of transcriptomic change, the recurrence of similar patterns across multiple tissues and independently-supported pathways that parallel mammalian hibernation seem to reflect that we have identified genuine features of hummingbird torpor biology. There are not only clear differences in the molecular players implicated, but also the scale of reduction of core body temperature and metabolic rates. Future integration of different-omics tools alongside functional experiments examining effects of such regulation will be essential for understanding how these birds enter and survive these states. With more avian torpor studies at the molecular scale, we can begin to identify the evolutionary and physiological mechanisms underlying these drastic 60-90% reductions in metabolic rate, and sometimes >20°C drops in body temperature, that these vertebrates are able to sustain for many hours a night.

## Materials and Methods

### Study animals and housing

Male Anna’s hummingbirds (*Calypte anna*, n=18) were captured at the George Fox University campus in Newberg, Oregon between June and July 2021 in the non-breeding season using modified Hall traps at nectar feeders. Birds were weighed, housed for <24 hours, and fed prior to the experiment with 20% nectar solution if measured the same night, or half-strength nectar with NektarPlus if measured the next night. For each bird, the final feeding time was between 19:30–21:15 h, after which it was weighed and placed into the experimental chamber for acclimation.

### Respirometry and thermal imaging

We used respirometry to estimate metabolic rates, and thermal imaging to measure surface temperatures of birds. Experimental chambers were designed to allow for simultaneous open-flow push respirometry and thermal video recordings through a DLC-coated silicon window. We used a Field Metabolic System (FMS, Sable Systems) to record metabolic rates following published protocols (2). Flow rates through the chamber were maintained at 800–840 ml/min using a Mass Flow System (Sable Systems) and sub-sampled by the FMS at 370 ml/min. Data on oxygen consumption, carbon dioxide production, and water vapour content of the air from the chambers were collected at 4 Hz to estimate metabolic rates. Warthog LabHelper software was used to collect data, and Warthog LabAnalyst to analyse them (<u>warthog.ucr.edu</u>). We used a factory-calibrated FLIR SC6701 infrared video camera to watch the birds live and record thermal videos for 10 seconds every 5 minutes using FLIR ResearchIR software (FLIR Systems, Inc., Wilsonville, OR, USA). We extracted maximum bird temperature (usually eye or shoulder) and background temperature (representing air temperature from a representative frame from each video) following published methods (67). We also recorded temperature data from the outside air and from the chamber using Type-T thermocouples and a TC-2000 (correlation of background temperature from IR camera with ambient temperature thermocouple adj R^2^ = 0.9972, p < 2.2e-16).

### Physiological state classification and tissue collection

Normothermic birds were those that maintained consistently high surface body temperatures in the sampling period (30–36°C). Transition birds were sampled halfway between their normothermic temperature and ambient temperature, in the middle of their entry into torpor (22–27°C). Birds in deep torpor (15–21°C) were sampled once they maintained their surface body temperatures within 3°C of ambient temperature for 10 minutes (Figure 1). Six birds in each state were sacrificed using an isoflurane-soaked cotton ball placed in the air flow path to the chamber. Birds were removed within one minute post-treatment and all tissues of interest (proximal gut, medial gut, distal gut, lungs, heart, liver and pectoral muscle) were extracted and preserved in RNALater within 15 minutes from isoflurane start time. Tissues were stored in RNALater at 4°C for 24 hours, after which excess RNALater was removed and samples were stored at −80°C until RNA extraction. To account for circadian regulation of gene expression, birds in different physiological states were sampled in a restricted time window between 2300 h and 0200 h. Four birds were sampled outside of this window at 22:37, 02:03, 02:14 and 02:40, with two of these in transition and one bird each in normothermy and torpor. Given that torpor entry in Anna’s hummingbirds (*Calypte anna*) can take between 1 and 2.5 hours (Figure S1A, S1B), moderate-to large-scale active mRNA turnover is biologically probable by torpor onset.

### RNA extraction, library preparation and sequencing

RNA was isolated using a modified Trizol protocol and sample quality checks were performed using spectrophotometry. Poly(A)+ mRNA selection and stranded libraries were generated using the NEBNext Poly(A) mRNA Magnetic Isolation Module and the NEBNext Ultra II Directional RNA Library Prep Kit (New England Biolabs) with 1 μg of total RNA input. Libraries were TruSeq-barcoded, quantified via Qubit (dsDNA HS kit; Thermo Fisher Scientific), and assessed for size distribution on an Agilent Fragment Analyzer prior to pooling. Sequencing was performed on an Illumina NovaSeq6000 platform to yield 2 × 150 bp paired-end reads, targeting a minimum depth of 20 million reads per library. This generated an average of 28 million raw reads (∼30 GB of raw data) per sample.

### Bioinformatic analyses

#### Read trimming and alignment

After quality filtering using TrimGalore under a stringent paired setting, an average of 25 million reads remained (Figure S2A). We used STAR (68) to index the *Calypte anna* reference genome (NCBI genome assembly: bCalAnn1_v1.p) against its GTF annotation and map trimmed reads. Of the 16230 genes on the reference, nearly 22% were uncharacterised “LOC” genes. Our mapping rates averaged 90%, with an average of 85% of reads mapping uniquely to a particular gene (Figure S2B).

#### Modelling expression of temperature-responsive genes

Gene expression could shift before torpor entry, early/late in torpor entry, or linearly as the bird’s body temperature drops. To assess which genes showed a temperature-specific or state-specific response, and whether they changed expression linearly or step-wise, we ran a series of models on the full gene expression matrix. The rlog-transformed expression matrix was first standardized to Z-scores within each tissue to account for differences in baseline expression of genes across tissue types. We first identified responsive genes by comparing an intercept-only linear mixed-effects model against a model with a model containing surface temperature (continuous, scaled to mean 0 and SD 1) and physiological state (a three-level factor: normothermy, transition, torpor), using a likelihood-ratio test with Benjamini–Hochberg correction across genes (p-adj < 0.05) and bird ID as random intercept in both models (multiple tissue samples from the same individual). For responsive genes, we compared four models by AIC: a linear model for surface temperature; an early-step model (normothermy vs transition and torpor); a late-step model (normothermy and transition vs torpor); and a state-means model (a separate mean for each of the three states). Genes were classified as *linear* when only the linear model was supported, *non-linear* when it was not, and *indistinguishable* when the linear model and a non-linear model were both supported (ΔAIC of within 2 units). *Non-linear* genes were further labelled *early step*, *late step*, or *graded/non-monotonic*; the last label denotes genes for which only the state-means model was supported, i.e., a pattern that neither a line nor a single step capture. To test for continuous temperature dependence beyond state, we compared the state-means model with a model that added temperature. Because screening with a temperature term can inflate this effect, we calibrated it against a permutation null (N=100). In each permutation, (i) temperature values were shuffled among birds within each state (with all tissues within each bird), which preserves state differences but removes any temperature effect beyond state; (ii) the screening step was repeated on the shuffled data; and (iii) the genes that passed the screen were classified, and tested for a temperature effect beyond state, exactly as in the observed data. To save computing time, the screen in each permutation was run only on genes with p < 0.05 in the observed screen, because genes above this threshold are very unlikely to pass once temperature is randomised; Benjamini–Hochberg correction still used the original number of tests across all genes. To test whether the responses track metabolic rate rather than surface temperature, we repeated the full analysis, including the permutation null, with log₁₀ mean VO₂ (averaged over the same ten-minute window as surface temperature) in place of surface temperature as the continuous predictor (Figure S18).

#### Differential gene expression analyses

To identify which genes were differentially expressed across metabolic states, raw reads were first filtered to retain only features with a total count sum >10 across all samples and rlog-stabilised. Principal Component Analysis (PCA) and Euclidean distance heatmaps identified three outlier samples that were removed from downstream analyses (Figure S2C, S2D). An additive model framework (design = ∼ Tissue + Metabolic State) was used to examine the effects of heterothermy across tissue types. To extract tissue-specific responses without subsetting by tissue type, a grouped factor design variable was used (e.g., Proximal Gut Deep Torpor). A generalised linear model was fitted to the entire dataset using DESeq2 (design = ∼ Group). Three pairwise contrasts were evaluated for each tissue: (i) Normothermy vs Deep Torpor, (ii) Normothermy vs Transition, and (iii) Transition vs Deep Torpor. Transcripts were classified as significantly differentially expressed genes (DEGs) if they met a Benjamini–Hochberg false discovery rate threshold of p-adjusted < 0.05 combined with a shrinkage-adjusted absolute effect size threshold of log2FC > 0.58.

#### Weighted Gene Correlation Network Analysis (WGCNA)

Weighted Gene Correlation Network Analysis (WGCNA) was employed to identify clusters of genes (modules) that show similar expression patterns across samples. WGCNA networks were constructed independently for each tissue type. Network construction was carried out with a block size of 15000 on the signed topological overlap matrix with a minModuleSize of 20 and mergeCutHeight of 0.3 after raising it to the appropriate power. The dynamic tree cut algorithm followed by merging of highly correlated eigengenes resulted in stable module definitions (Figure S4). To examine correlations between module eigengenes and physiological traits of interest, we used multiple regression with the lme4 package to evaluate the independent effects of metabolic rate and surface body temperature following recent methodological advancements (28) (Figure S16). Hub genes of the retained modules were identified by module membership (kME; Figure S17). Modules that were also independently correlated with time of the night were excluded from the pool of shortlisted modules, as these likely reflect circadian responses rather than primarily the torpor entry process.

#### Functional annotation and KEGG enrichment

Since the *Calypte anna* genome lacks KEGG annotations, protein-coding sequences were functionally annotated using the KEGG Automatic Annotation Server (KAAS). Using this approach, 76.5% of protein isoforms and 83.7% of genes in the genome were assigned at least one KEGG Orthology identifiers (KO). In total, 9,945 unique KOs were recovered, with 12,399 gene–KO associations. KEGG enrichments were performed on hits obtained from two complementary pipelines: (i) DEG analyses and (ii) WGCNA modules whose eigengenes correlated with physiological traits. KEGG enrichment analyses were conducted using clusterProfiler with a genome-restricted KO background comprising all KOs observed in the annotated gene set.

#### Targeted pathway-level analysis

To test whether specific biological programmes shift in torpor, we analysed pre-specified gene sets for eleven target pathways (TCA cycle, oxidative phosphorylation, PPAR signalling, thermogenesis, circadian rhythm, spliceosome, translation, transcription, cell cycle, PI3K–Akt signalling and FoxO signalling). For each pathway, we took the whole KEGG pathway and searched for each of its human gene symbols in our annotation; genes with a matching symbol (“matched” genes) were included, and genes without one were not tested. We also defined curated functional groups from gene function before we inspected results. A gene set (“set”) is the unit of analysis in the targeted tests and is either a KEGG set (the whole KEGG pathway) or a curated set (a functional group of genes within that pathway, for example mitochondrial translation within translation). We use these three terms throughout: target pathway (one of the eleven), KEGG set and curated set. In total, 87 sets (KEGG and curated) across the eleven pathways were tested, and tissues with fewer than three birds in a state (lungs) were excluded. We compared normothermic with deep-torpor birds, with bird identity as a random effect because tissues from the same bird are not independent. For each set we asked two questions: whether the set shifts more than other genes (competitive test, camera) (68) and whether it shifts at all (self-contained test, based on the mean Z-score of the set), each time per tissue and across tissues combined (“pooled”). As a direction-agnostic check, mroast asks whether a set’s genes shift in either direction. Effect sizes are Hedges’ g (torpor minus normothermy, in units of within-group variability), and p-values were Benjamini–Hochberg-adjusted within each pathway. Throughout the Results, we report camera first, then the self-contained test, then mroast where relevant, and we name the test before each value; pooled results come before per-tissue results. We report adjusted p-values (p-adj) throughout; unadjusted values are labelled “p-nominal”. More details of the structure of these tests are in the Supplementary Methods.

## Data Availability

Data are available as an attached.zip file. All code is on Github at https://github.com/Notarcha/Hummers_Torpor_-Transcriptomics_Code.

## Ethics Statement

This work was done under the following US Fish and Wildlife and Animal Ethics permits. USFWS - MB93258B (2021), MB93258B (2022). Oregon: 039-21 (2021), 039-22 (2022). Arizona: SP407009 (2021), SP803179 (2022). George Fox University IACUC# - 010.

## Acknowledgments

The authors thank Bronwyn Butcher for training AS in RNA extraction protocols and providing extensive laboratory support. We are deeply grateful to Irby Lovette for his generosity and support, as well as the members of the Lovette Lab—specifically Sabrina McNew and Gemma Clucas—for their invaluable feedback on study design. We thank Ann Tate for help with RNA extractions; Jen Grenier and Faraz Ahmed for sequencing and preliminary RNA-seq data processing; the Cornell BioHPC for excellent computational facilities; and Jeff Glaubitz and Qi Sun for training and analytical support. Administrative assistance was kindly provided by Teresa Arnold at George Fox University. We thank Anagha Mohan for doing preliminary analyses and giving the work some direction. We also thank Nora Prior, Maren Vitousek and the Vitousek Lab, Hans Hoffman, Brian Barnes, Praveen Sethupathy, Sinisa Hrvatin, Manish Jaiswal, Ullas Kolthur-Seetharam, Aprotim Mazumder, Mrinal Srivastava, Jay Phadke, and Keshav Srinivasan for discussions on the work presented here. We thank Sophia Wolfe and Shenni Liang for comments on the manuscript draft and discussions. This work was supported by the Cornell Biotechnology Resource Center (BRC). Funding was provided by the Rose Postdoctoral Fellowship (to AS); National Geographic (Grants NGS-91718R-21 and EC-53404R-21); the Center for Vertebrate Genomics (CVG Scholar Program, to AS); a postdoctoral research grant from the American Ornithological Society (AOS, to AS); the Tata Institute of Fundamental Research Hyderabad (TIFRH); and the Department of Atomic Energy (DAE) under Project Identification No. RTI 40007 to AS and HK.

