## Supplementary Methods, Figures, Tables for "Hummingbird torpor recapitulates key molecular signatures of hibernation without large-scale transcriptomic remodelling"

#### Animal chamber design

Chambers were 10.16cm (inner dimensions) acrylic cubes. All sides except the front were opaque white to help birds settle easily; the front side was clear acrylic to allow light through. The front face had a circular opening in which we fit a 76 mm diameter DLC-coated silicon window (Edmund Optics) for thermal imaging. The top was removable for placement of the bird in the chamber and sealable with a clamp.

#### RNA extraction

After the first phase separation, an additional chloroform extraction step of the aqueous layer was performed in Phase-lock Gel heavy tubes (Quanta Biosciences), followed by addition of 1 µl GlycoBlue (Thermo Fisher) immediately prior to isopropanol precipitation and three washes of the RNA pellet with 75% ethanol.

#### STAR alignment and count generation

STAR was run with the flags --outFilterIntronMotifs RemoveNoncanonical, --outSAMtype BAM SortedByCoordinate, and --quantMode GeneCounts, while retaining all other parameters at their defaults. The resulting read counts for each sample were pooled into a comprehensive master count matrix by selecting the reverse-stranded column from the STAR gene-count output, matching the strand-specific architecture of our library preparation.

#### WGCNA pre-processing and network construction

Prior to network analysis, outliers from the read count matrix were removed and the matrix was rlog-stabilized and subset by tissue type to examine outlier genes using the goodSamplesGenes function of the WGCNA package (Langfelder & Horvath, 2008). To reduce noise prior to WGCNA, genes with low expression values were filtered out and only the 80% most variable genes were retained. A soft-power thresholding simulation was performed to identify the power that best approximated a scale-free network while maintaining moderate levels of connectivity. In all cases, networks approached scale-free topology ($R^2 > 0.80$) across different powers for each tissue type (Figure S4).

#### KEGG annotation of the *Calypte anna* genome

All predicted protein isoforms from the reference genome were submitted to KAAS, and KEGG Orthology (KO) identifiers were assigned at the protein level using the bidirectional best hit (BBH) method against a curated panel of avian KEGG reference genomes. These included *Gallus gallus* (gga), *Taeniopygia guttata* (tgu), *Anas platyrhynchos* (ana), *Meleagris gallopavo* (mel), *Columba livia* (cot), *Falco peregrinus* (fpg), *Apteryx australis* (asg), *Calypte anna* (cal), *Aptenodytes forsteri* (apt), and *Charadrius vociferus* (chr). Protein-level KO annotations were subsequently collapsed to the gene level by retaining unique gene–KO mappings.

#### Targeted pathway-level analysis

#### Supplementary Methods

*Gene sets.* To test whether specific biological programmes shift in torpor, we analysed pre-specified gene sets for eleven target pathways (TCA cycle, oxidative phosphorylation, PPAR signalling, thermogenesis, circadian rhythm, spliceosome, translation, transcription, cell cycle, PI3K–Akt signalling and FoxO signalling). Each target pathway comprised the whole KEGG pathway, whose human gene symbols we searched for in our annotation ("matched" genes are those found), and curated functional groups defined from gene function before we inspected results. A gene set ("set") is the unit of analysis in the targeted tests and is either a KEGG set (the whole KEGG pathway) or a curated set (a functional group of genes within that pathway, for example mitochondrial translation within translation). We use these three terms throughout: target pathway (one of the eleven), KEGG set and curated set. Genes absent from the dataset were not tested, nor were sets with fewer than three genes present; in total, 87 sets (KEGG and curated) across the eleven pathways were tested, and tissues with fewer than three birds in a state (i.e., lungs) were excluded. Genes that belong only to the whole KEGG pathway (and not to any curated set) contribute to the KEGG set test but were not tested or reported individually.

*Expression data and standardisation.* All analyses started from the rlog-normalised expression matrix. For the self-contained tests, each gene's rlog expression was standardised to a Z-score within each tissue (the tissue mean was subtracted and the result divided by the standard deviation across that tissue's normothermic and deep-torpor samples), so that genes and tissues with different baseline expression are comparable.

*Gene-level tests.* Normothermy was compared with deep torpor using per-tissue Z-scores, and bird identity was modelled as a random effect so that tissues from the same bird were not treated as independent. For each gene we obtained a test statistic per tissue and with the tissues combined: per tissue, a moderated t-statistic (limma) on rlog expression; combined, a model with tissue and state terms, a bird random intercept (duplicateCorrelation) and unequal variances per tissue × state. "Pooled" refers to all tissues combined in this way, and "per tissue" to a separate test within one tissue.

*Set-level tests.* For each set we asked two questions: whether the set shifts more than other genes (competitive test, camera) and whether it shifts at all (self-contained test, based on the mean Z-score of the set), each time per tissue and across tissues combined. The competitive test used rlog expression with the inter-gene correlation estimated for each set; for the combined test, the data were transformed to account for the same bird correlation and variance structure, so bird identity was also modelled in camera. The self-contained test used the per-sample set score (the mean Z-score of the set's genes) with Welch's t-test within each tissue and, across tissues combined, a linear mixed model with bird as a random intercept and unequal variances per tissue × state (likelihood-ratio test). As a direction-agnostic check, we also ran mroast (the "mixed" statistic, 999 rotations) on the same data and model as camera, which asks whether the genes of a set shift in either direction.

*Effect sizes and multiple testing.* Effect sizes are reported as Hedges' g, the difference between torpor and normothermy expressed in units of the within-group variability (a signal-to-noise measure, where values near ±1 mean the shift is about as large as the spread between individuals). P-values were corrected for multiple testing (Benjamini–Hochberg), separately within each pathway: across all sets (combined tests) or set × tissue tests (per tissue), and across all genes (combined) or gene × tissue tests (per tissue), so that a gene present in more than one pathway can have a different p-adj in each.

*Reporting conventions.* Throughout the Results, we report camera first, then the self-contained test, then mroast where relevant, and we name the test before each value; pooled results come before per-tissue results. We report adjusted p-values (p-adj) throughout; unadjusted values are labelled “p-nominal”.

### Supplementary Figures and Tables

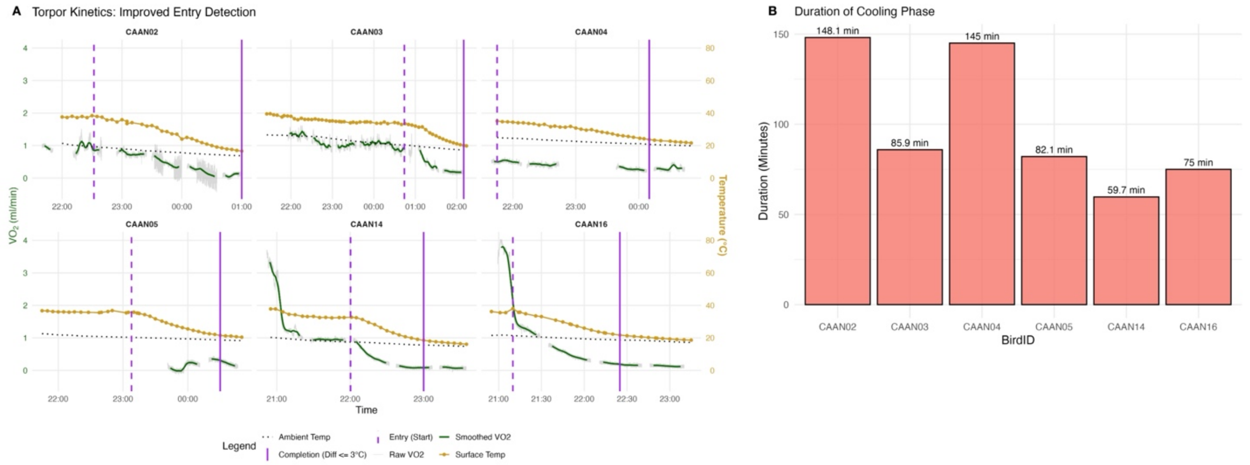


Figure S1: A - Dual-axis plots display traces for oxygen consumption ($VO_2$, left axis) and surface temperature ($T_s$, right axis) for individuals during the cooling phase. The onset of torpor entry (dashed vertical purple line) was identified by the maximal negative slope of the surface temperature decline (orange trace). The completion of entry (solid vertical purple line) was defined as the point where $T_s$ equilibrated to within 3°C of the ambient temperature (black dotted line). Raw $VO_2$ data (grey traces) are overlaid with smoothed $VO_2$ measurements (green traces). Gaps in the $VO_2$ traces indicate timepoints excluded due to baseline adjustments or technical artifacts (e.g., sensor issues). B - Timescales of torpor entry in each individual of *Calypte anna*.

### Supplementary Figures and Tables

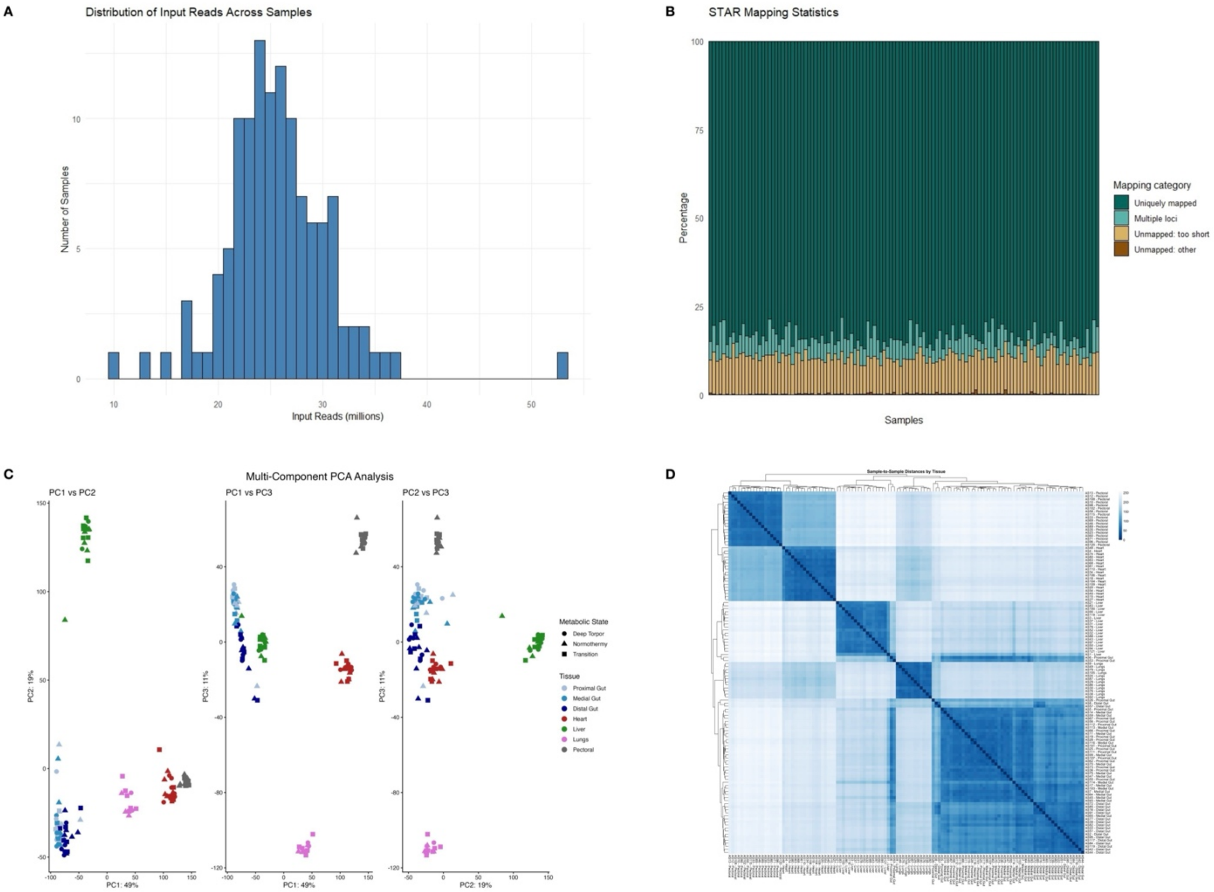


Figure S2: A - Distribution of number of reads available for mapping with STAR post quality filtering using TrimGalore. B - Mapping rates of trimmed reads from different samples of *Calypte anna* to its reference genome (GCF_003957555.1). C - Multicomponent PCA depicting separation between transcriptomic profiles of different tissue types and physiological states. D - Pairwise Euclidean distance of samples across tissue types to identify outliers.

### Supplementary Figures and Tables

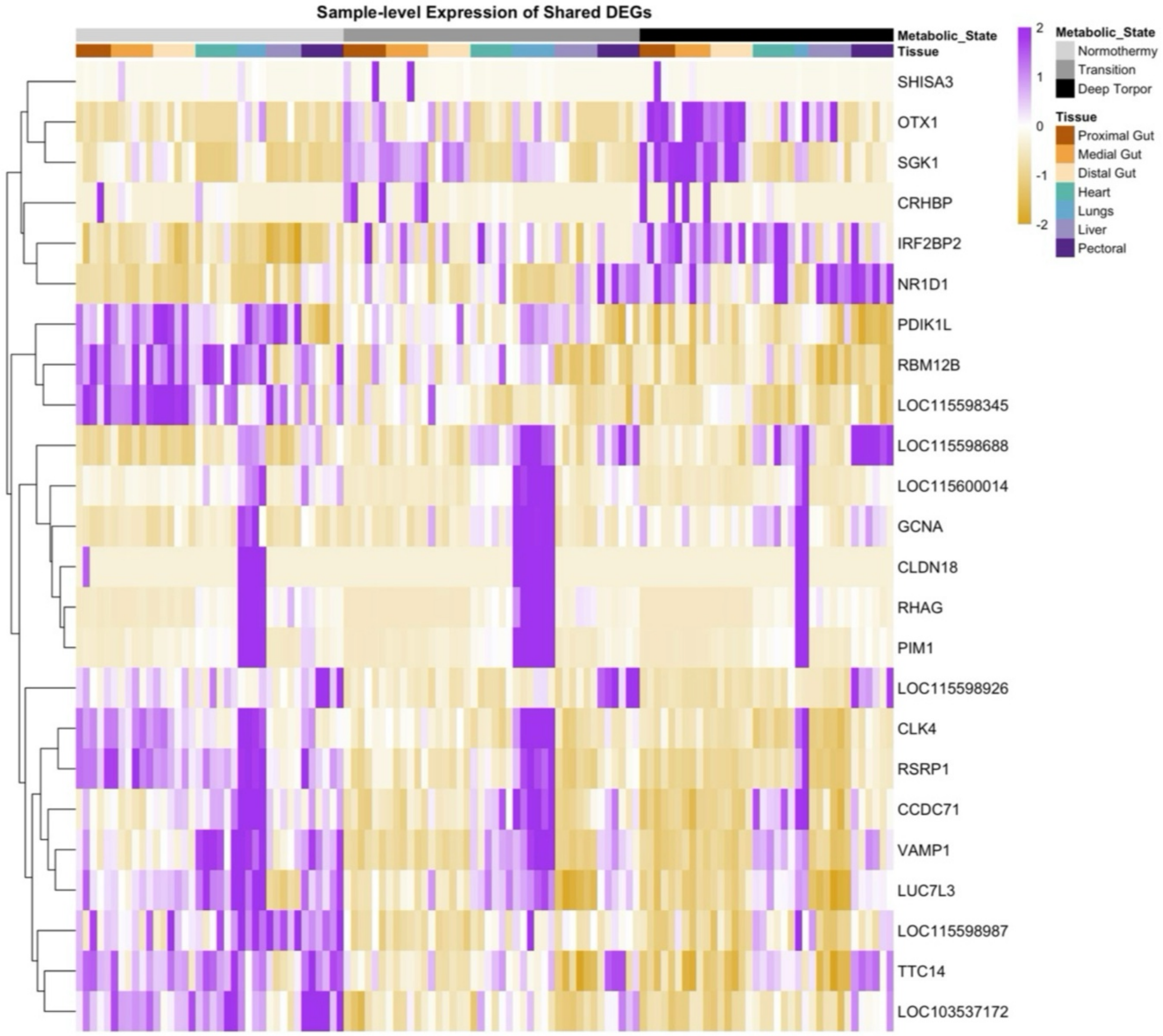


Figure S3: Sample-to-sample variation in differential expression of shared DEGs (expression in torpor - expression in normothermy) across tissue types. Each row corresponds to a DEG and each column corresponds to a sample from a given tissue type.

### Supplementary Figures and Tables

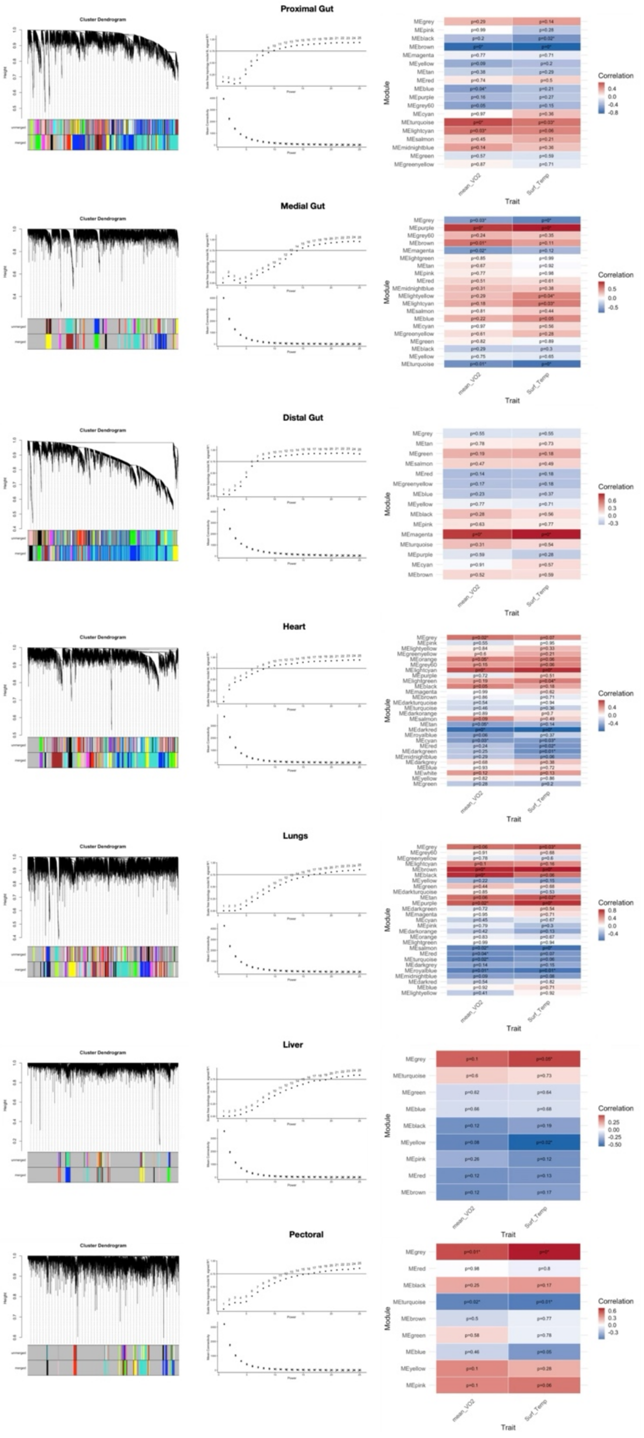


Figure S4: Column 1 - Cluster dendrograms depicting merging of modules and dynamic cut-tree algorithm effectively merging gene members into modules. Column 2 - Soft thresholding algorithms depicting how gene regulatory networks for different tissue types approach scale free-topology (a crucial assumption of WGCNA methods). Column 3 - Correlation maps of module eigen genes and how they correlate with physiological traits.

### Supplementary Figures and Tables

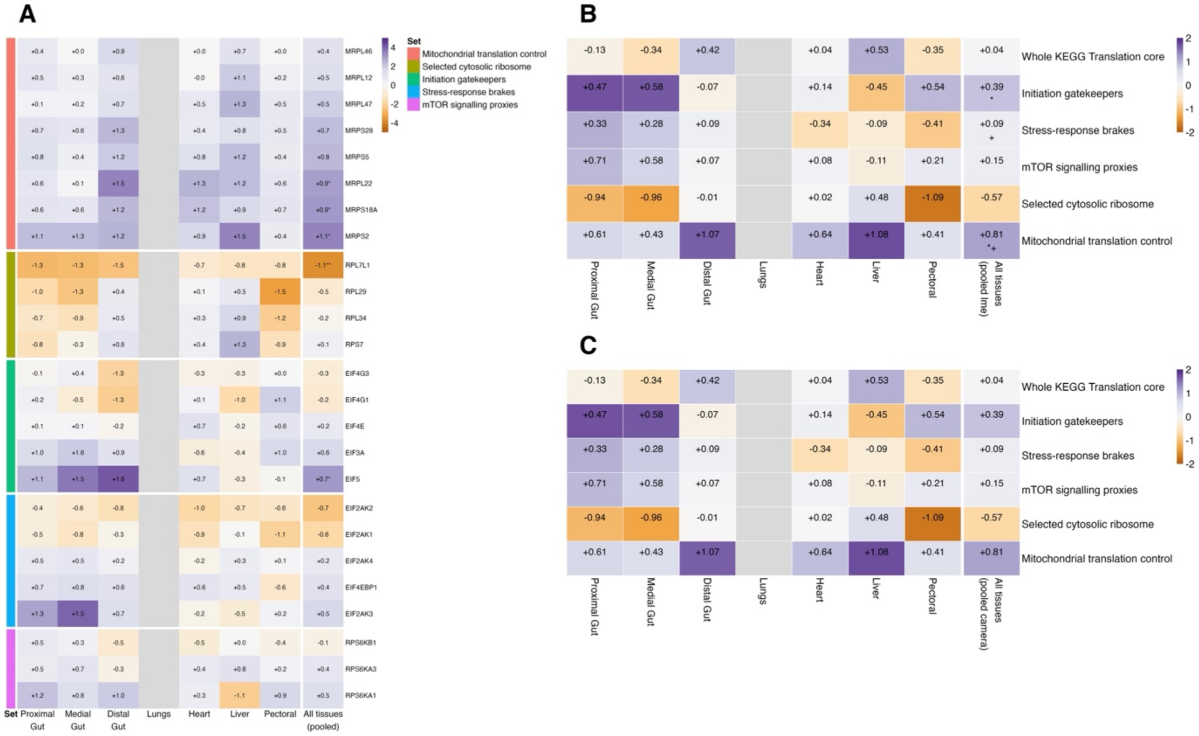


Figure S5: Mitochondrial translation and initiation factors are upregulated in torpor, whereas the cytosolic translation machinery remains unchanged. Panel A is a set of curated, interesting, genes from the sets in the legend; B and C compare the pathways using all the genes in that pathway. **(A)** Gene-level responses for curated translation genes, grouped by functional set. Colour shows the limma moderated $t$ for torpor versus normothermy (purple, higher in torpor; orange, lower; scaled to ±5). Numbers give the mean shift in per-tissue Z-score units (torpor − normothermy). Asterisks mark p-adj < 0.05 (*) and p-adj < 0.01 (**) (Benjamini–Hochberg), corrected across all gene × tissue tests for the tissue columns and across genes for the last column. **(B)** Self-contained tests of whether each set shifts at all. Colour shows Hedges' $g$ (torpor − normothermy; scaled to ±2) and numbers show the mean shift of the set in Z units. Asterisks mark p-adj < 0.05 (*) from Welch's $t$-test per tissue and from a mixed model for the pooled column. A plus sign (+) marks p-adj < 0.05 in the direction-agnostic mroast test. **(C)** Competitive tests (camera) with the same colour and number scheme as B. Grey cells were not tested (lungs: fewer than three birds in torpor).

### Supplementary Figures and Tables

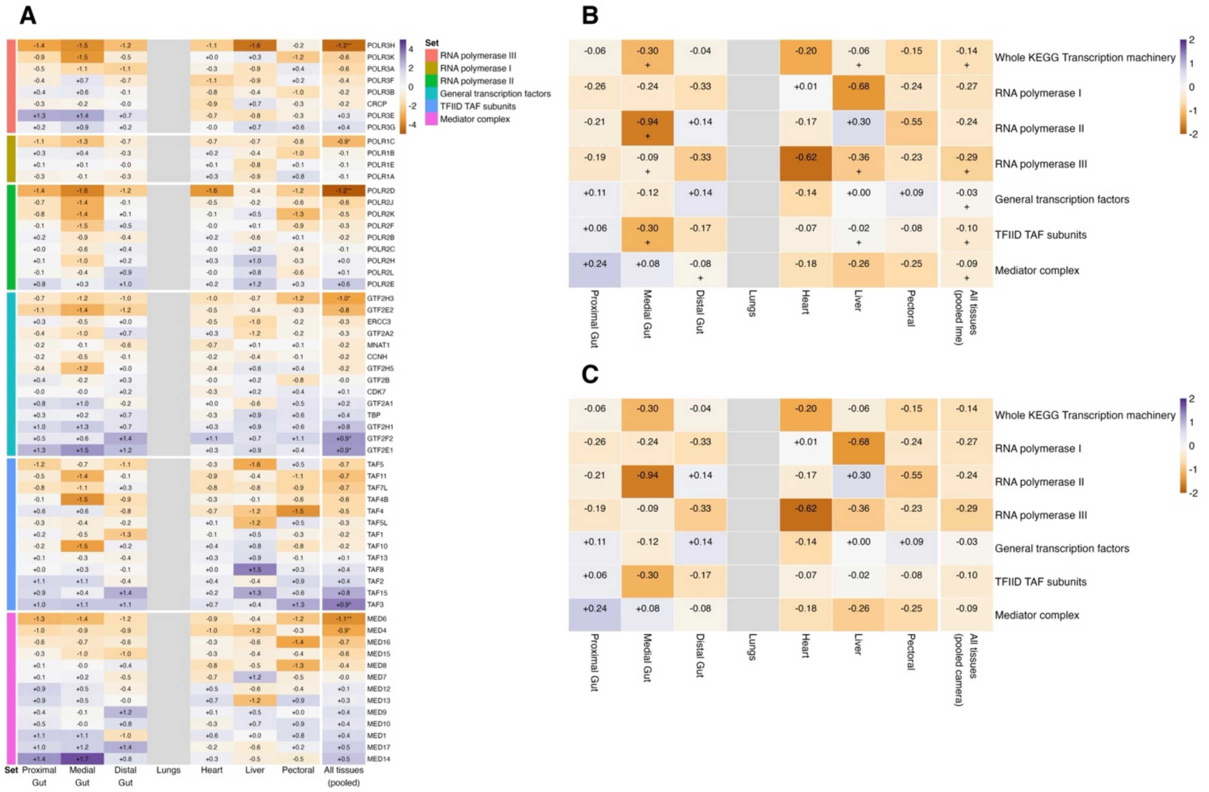


Figure S6: The transcription machinery as a whole is unchanged in torpor, although individual members shift in both directions. (A) Gene-level responses for genes in the tested sets, grouped by functional set. Colour shows the limma moderated t for torpor versus normothermy (purple, higher in torpor; orange, lower; scaled to ±5). Numbers give the mean shift in per-tissue Z-score units (torpor − normothermy). Asterisks mark p-adj < 0.05 (*) and p-adj < 0.01 (**) (Benjamini–Hochberg), corrected across all gene × tissue tests for the tissue columns and across genes for the last column. (B) Self-contained tests of whether each set shifts at all. Colour shows Hedges' g (torpor − normothermy; scaled to ±2) and numbers show the mean shift of the set in Z units. Asterisks mark p-adj < 0.05 (*) from Welch's t-test per tissue and from a mixed model for the pooled column. A plus sign (+) marks p-adj < 0.05 in the direction-agnostic mroast test. (C) Competitive tests (camera) with the same colour and number scheme as B. Grey cells were not tested (lungs: fewer than three birds in torpor).

### Supplementary Figures and Tables

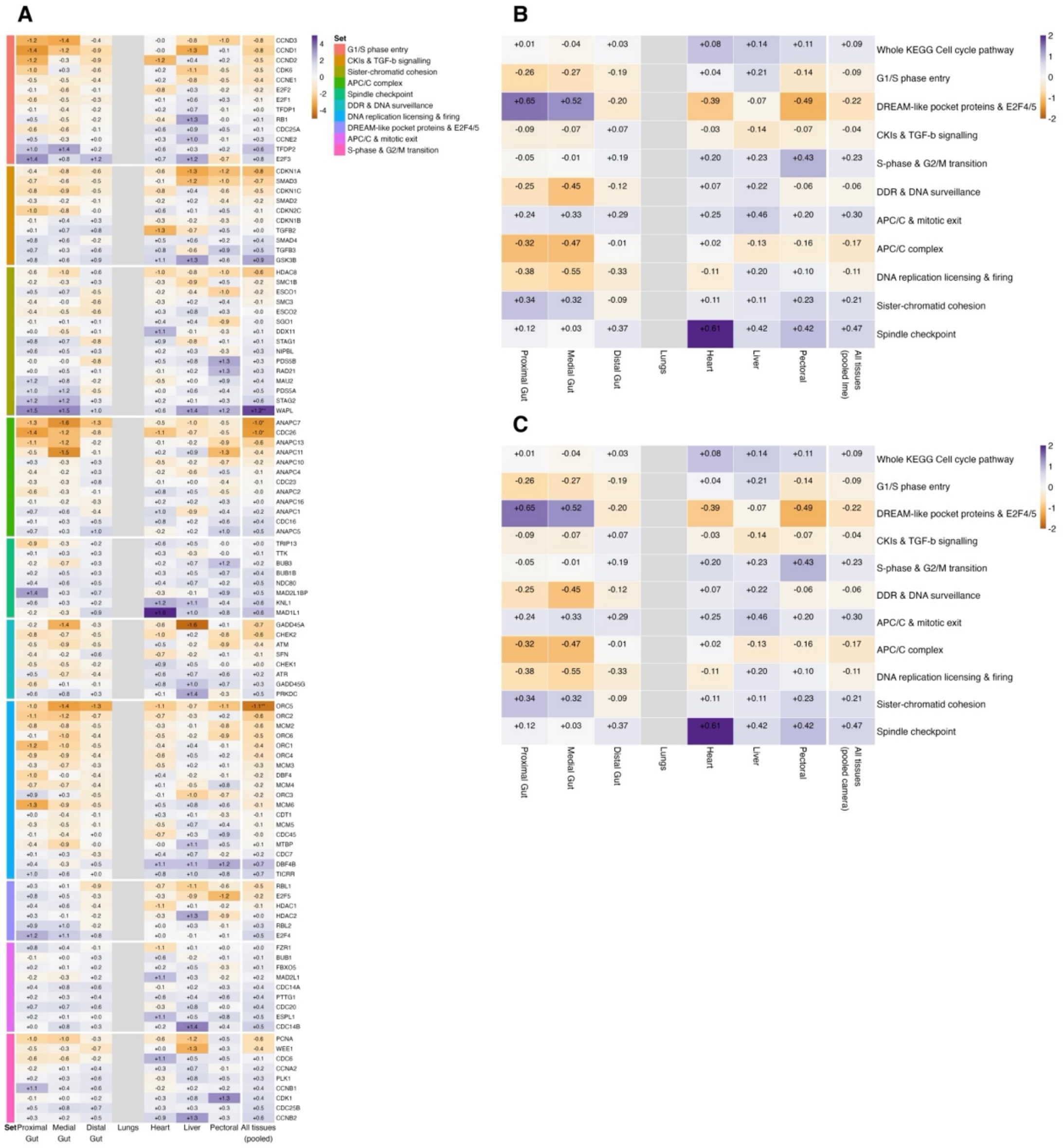


Figure S7: The cell-cycle machinery is unchanged in torpor, although individual members shift in both directions. (A) Gene-level responses for genes in the tested sets, grouped by functional set. Colour shows the limma moderated t for torpor versus normothermy (purple, higher in torpor; orange, lower; scaled to ±5). Numbers give the mean shift in per-tissue Z-score units (torpor − normothermy). Asterisks mark p-adj < 0.05 (*) and p-adj < 0.01 (**) (Benjamini–Hochberg), corrected across all gene × tissue tests for the tissue columns and across genes for the last column. (B) Self-contained tests of whether each set shifts at all. Colour shows Hedges' g (torpor − normothermy; scaled to ±2) and numbers show the mean shift of the set in Z units. Asterisks mark p-adj < 0.05 (*) from Welch's t-test per tissue and from a mixed model for the pooled column. A plus sign (+) marks p-adj < 0.05 in the direction-agnostic mroast test. (C) Competitive tests (camera) with the same colour and number scheme as B. Grey cells were not tested (lungs: fewer than three birds in torpor).

### Supplementary Figures and Tables

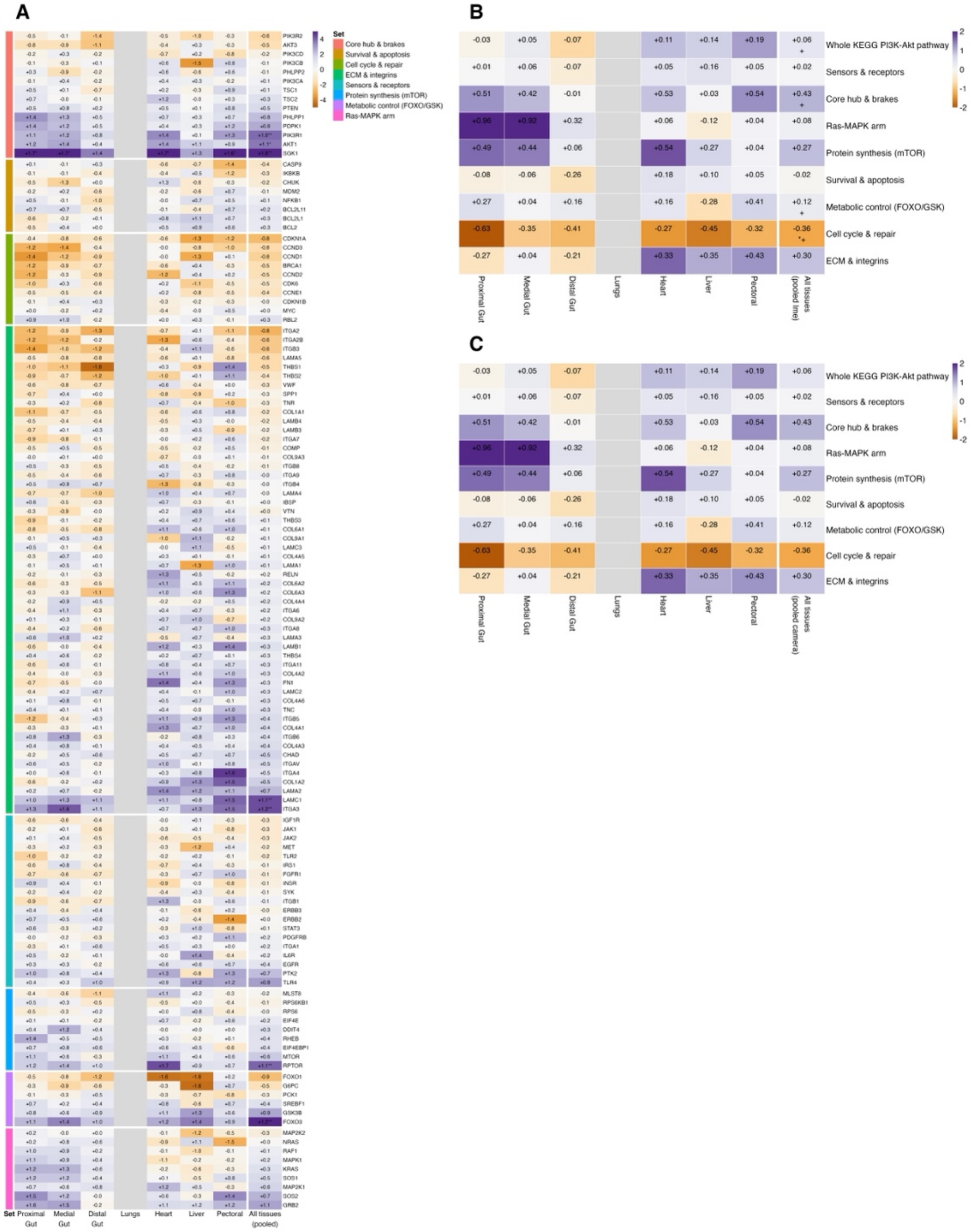

Figure S8: The Akt/SGK kinase module, led by SGK1, trends higher in torpor. (A) Gene-level responses for genes in the tested sets, grouped by functional set. Colour shows the limma moderated t for torpor versus normothermy (purple, higher in torpor; orange, lower; scaled to ±5). Numbers give the mean shift in per-tissue Z-score units (torpor − normothermy). Asterisks mark p-adj < 0.05 (*) and p-adj < 0.01 (**) (Benjamini–Hochberg), corrected across all gene × tissue tests for the tissue columns and across genes for the last column. (B) Self-contained tests of whether each set shifts at all. Colour shows Hedges' g (torpor − normothermy; scaled to ±2) and numbers show the mean shift of the set in Z units. Asterisks mark p-adj < 0.05 (*) from Welch's t-test per tissue and from a mixed model for the pooled column. A plus sign (+) marks p-adj < 0.05 in the direction-agnostic mroast test. (C) Competitive tests (camera) with the same colour and number scheme as B. Grey cells were not tested (lungs: fewer than three birds in torpor).

### Supplementary Figures and Tables

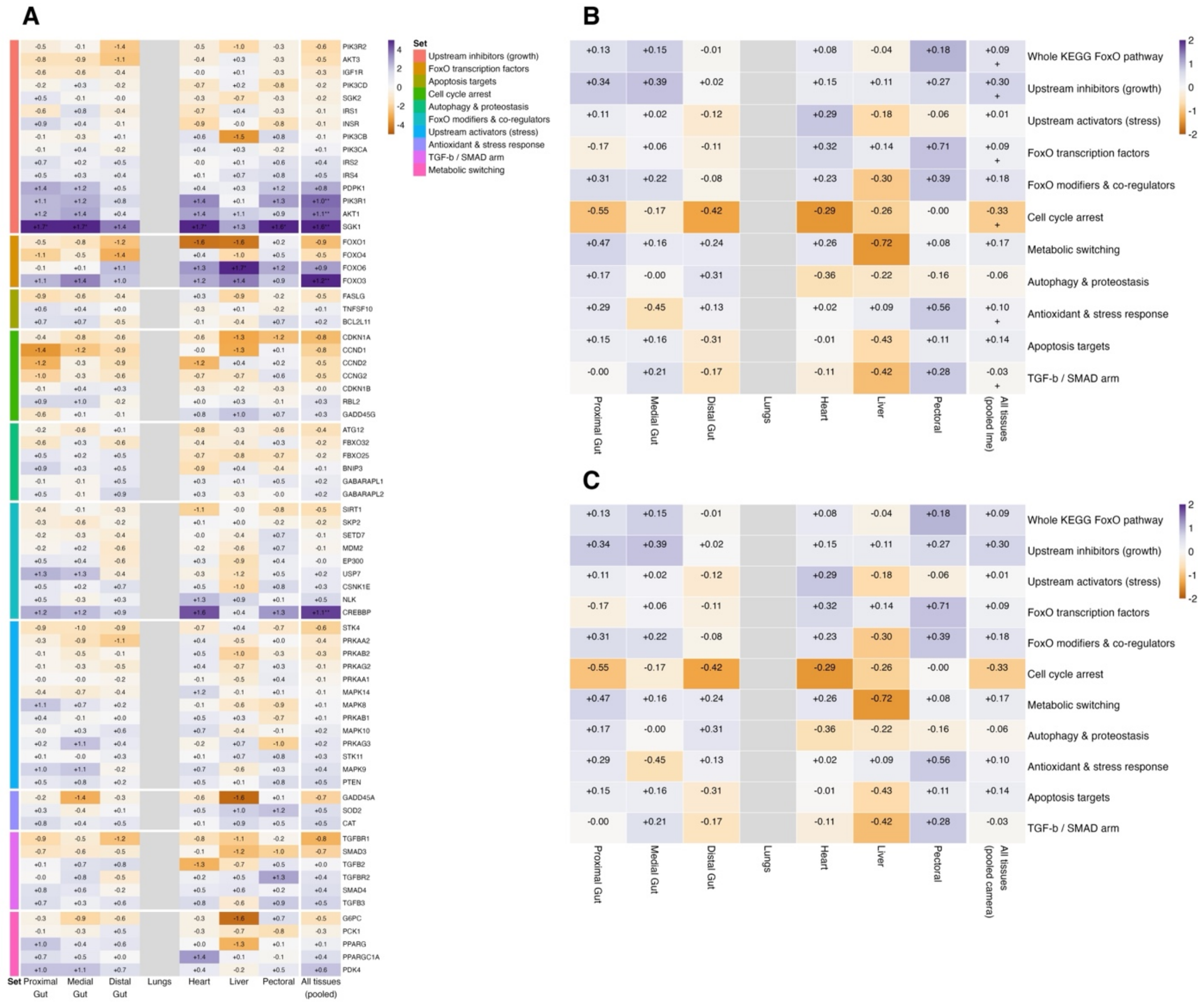


Figure S9: FoxO factors move in opposite directions in torpor, with FOXO3 higher and FOXO1 trending lower. (A) Gene-level responses for genes in the tested sets, grouped by functional set. Colour shows the limma moderated t for torpor versus normothermy (purple, higher in torpor; orange, lower; scaled to ±5). Numbers give the mean shift in per-tissue Z-score units (torpor − normothermy). Asterisks mark p-adj < 0.05 (*) and p-adj < 0.01 (**) (Benjamini–Hochberg), corrected across all gene × tissue tests for the tissue columns and across genes for the last column. (B) Self-contained tests of whether each set shifts at all. Colour shows Hedges' g (torpor − normothermy; scaled to ±2) and numbers show the mean shift of the set in Z units. Asterisks mark p-adj < 0.05 (*) from Welch's t-test per tissue and from a mixed model for the pooled column. A plus sign (+) marks p-adj < 0.05 in the direction-agnostic mroast test. (C) Competitive tests (camera) with the same colour and number scheme as B. Grey cells were not tested (lungs: fewer than three birds in torpor).

### Supplementary Figures and Tables

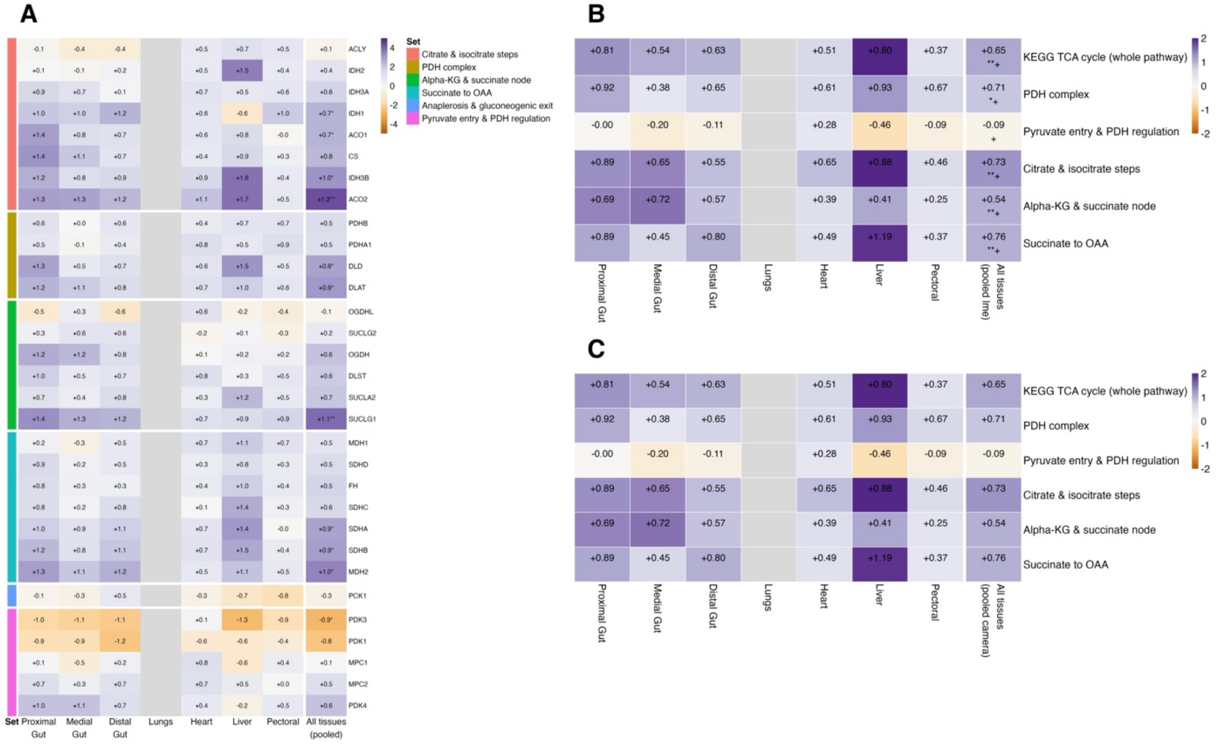


Figure S10: TCA cycle transcripts are modestly but consistently higher in torpor. (A) Gene-level responses for genes in the tested sets, grouped by functional set. Colour shows the limma moderated t for torpor versus normothermy (purple, higher in torpor; orange, lower; scaled to ±5). Numbers give the mean shift in per-tissue Z-score units (torpor − normothermy). Asterisks mark p-adj < 0.05 (*) and p-adj < 0.01 (**) (Benjamini–Hochberg), corrected across all gene × tissue tests for the tissue columns and across genes for the last column. (B) Self-contained tests of whether each set shifts at all. Colour shows Hedges' g (torpor − normothermy; scaled to ±2) and numbers show the mean shift of the set in Z units. Asterisks mark p-adj < 0.05 (*) from Welch's t-test per tissue and from a mixed model for the pooled column. A plus sign (+) marks p-adj < 0.05 in the direction-agnostic mroast test. (C) Competitive tests (camera) with the same colour and number scheme as B. Grey cells were not tested (lungs: fewer than three birds in torpor).

### Supplementary Figures and Tables

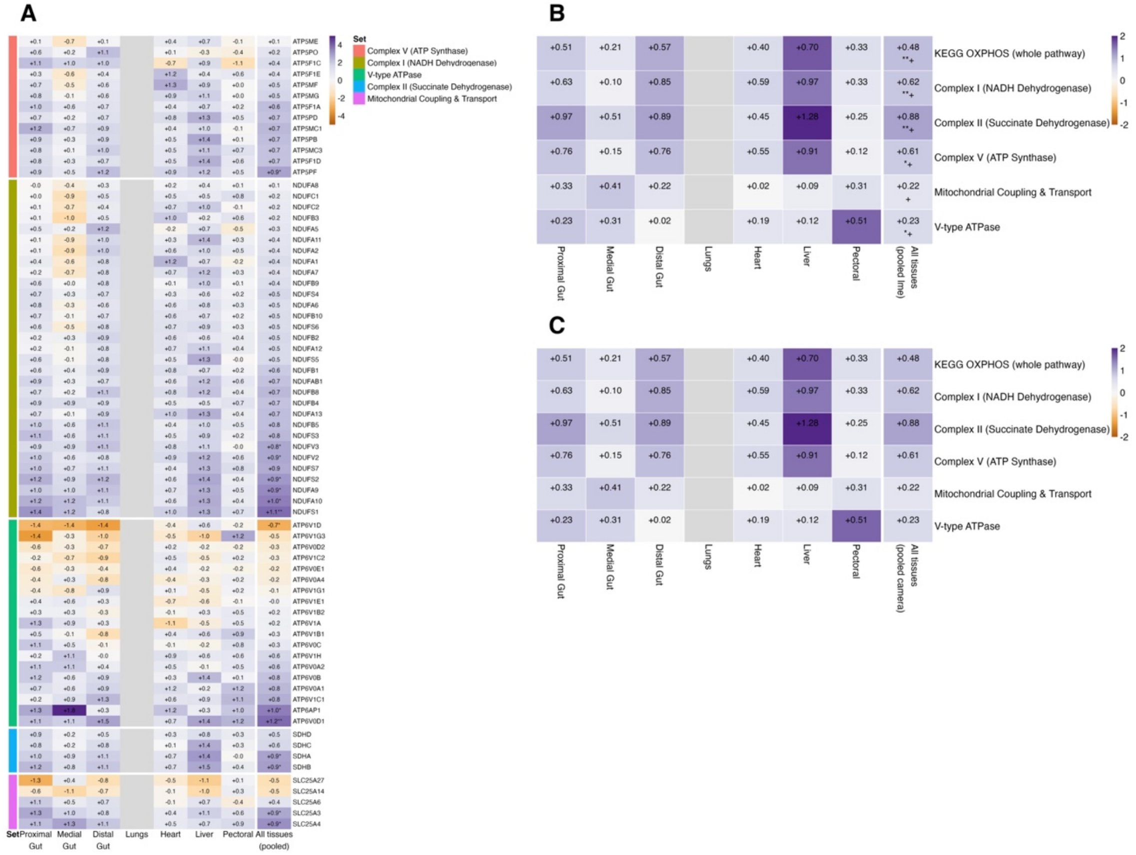


Figure S11: Oxidative phosphorylation transcripts show a modest, broad upward shift in torpor. (A) Gene-level responses for genes in the tested sets, grouped by functional set. Colour shows the limma moderated t for torpor versus normothermy (purple, higher in torpor; orange, lower; scaled to ±5). Numbers give the mean shift in per-tissue Z-score units (torpor − normothermy). Asterisks mark p-adj < 0.05 (*) and p-adj < 0.01 (**) (Benjamini–Hochberg), corrected across all gene × tissue tests for the tissue columns and across genes for the last column. (B) Self-contained tests of whether each set shifts at all. Colour shows Hedges' g (torpor − normothermy; scaled to ±2) and numbers show the mean shift of the set in Z units. Asterisks mark p-adj < 0.05 (*) from Welch's t-test per tissue and from a mixed model for the pooled column. A plus sign (+) marks p-adj < 0.05 in the direction-agnostic mroast test. (C) Competitive tests (camera) with the same colour and number scheme as B. Grey cells were not tested (lungs: fewer than three birds in torpor). Complexes III and IV and mitochondrially encoded subunits are absent from the expression matrix and are not shown.

### Supplementary Figures and Tables

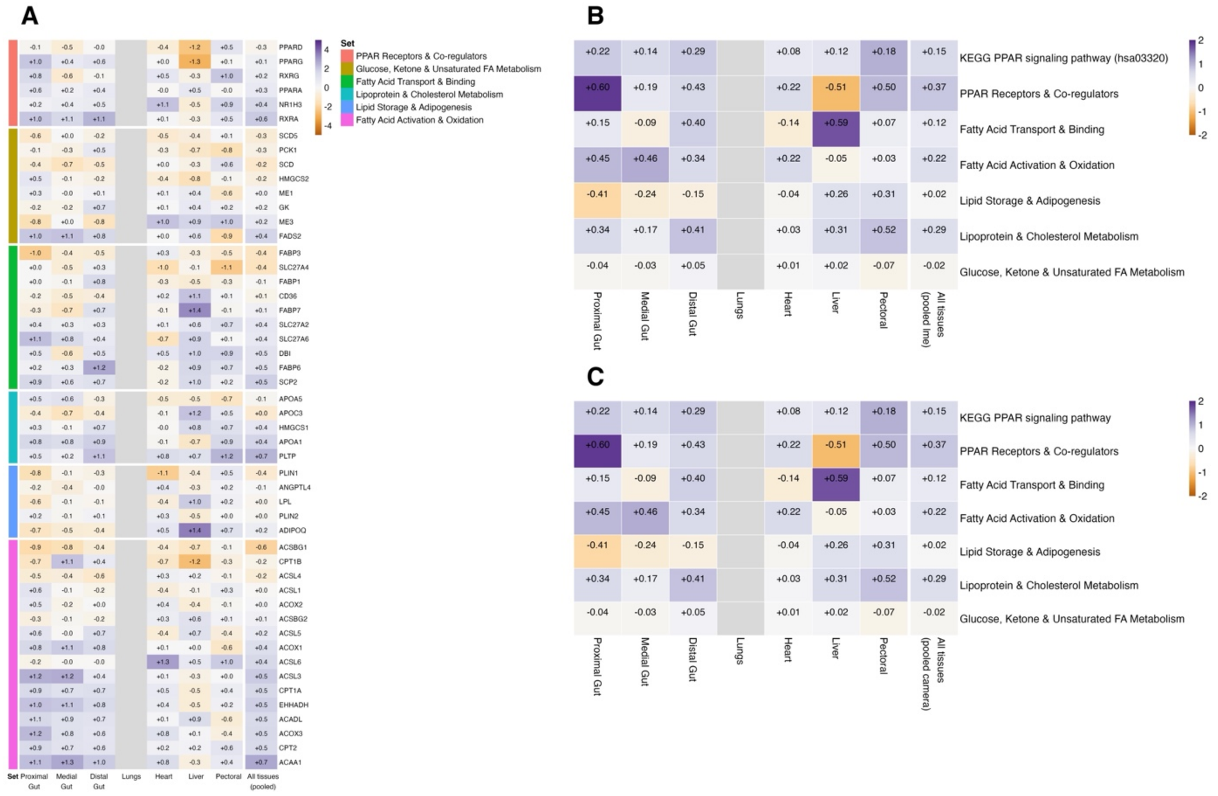

Figure S12: PPAR signalling shows no significant pathway-level change in torpor. (A) Gene-level responses for genes in the tested sets, grouped by functional set. Colour shows the limma moderated t for torpor versus normothermy (purple, higher in torpor; orange, lower; scaled to ±5). Numbers give the mean shift in per-tissue Z-score units (torpor − normothermy). Asterisks mark p-adj < 0.05 (*) and p-adj < 0.01 (**) (Benjamini–Hochberg), corrected across all gene × tissue tests for the tissue columns and across genes for the last column. (B) Self-contained tests of whether each set shifts at all. Colour shows Hedges' g (torpor − normothermy; scaled to ±2) and numbers show the mean shift of the set in Z units. Asterisks mark p-adj < 0.05 (*) from Welch's t-test per tissue and from a mixed model for the pooled column. A plus sign (+) marks p-adj < 0.05 in the direction-agnostic mroast test. (C) Competitive tests (camera) with the same colour and number scheme as B. Grey cells were not tested (lungs: fewer than three birds in torpor).

### Supplementary Figures and Tables

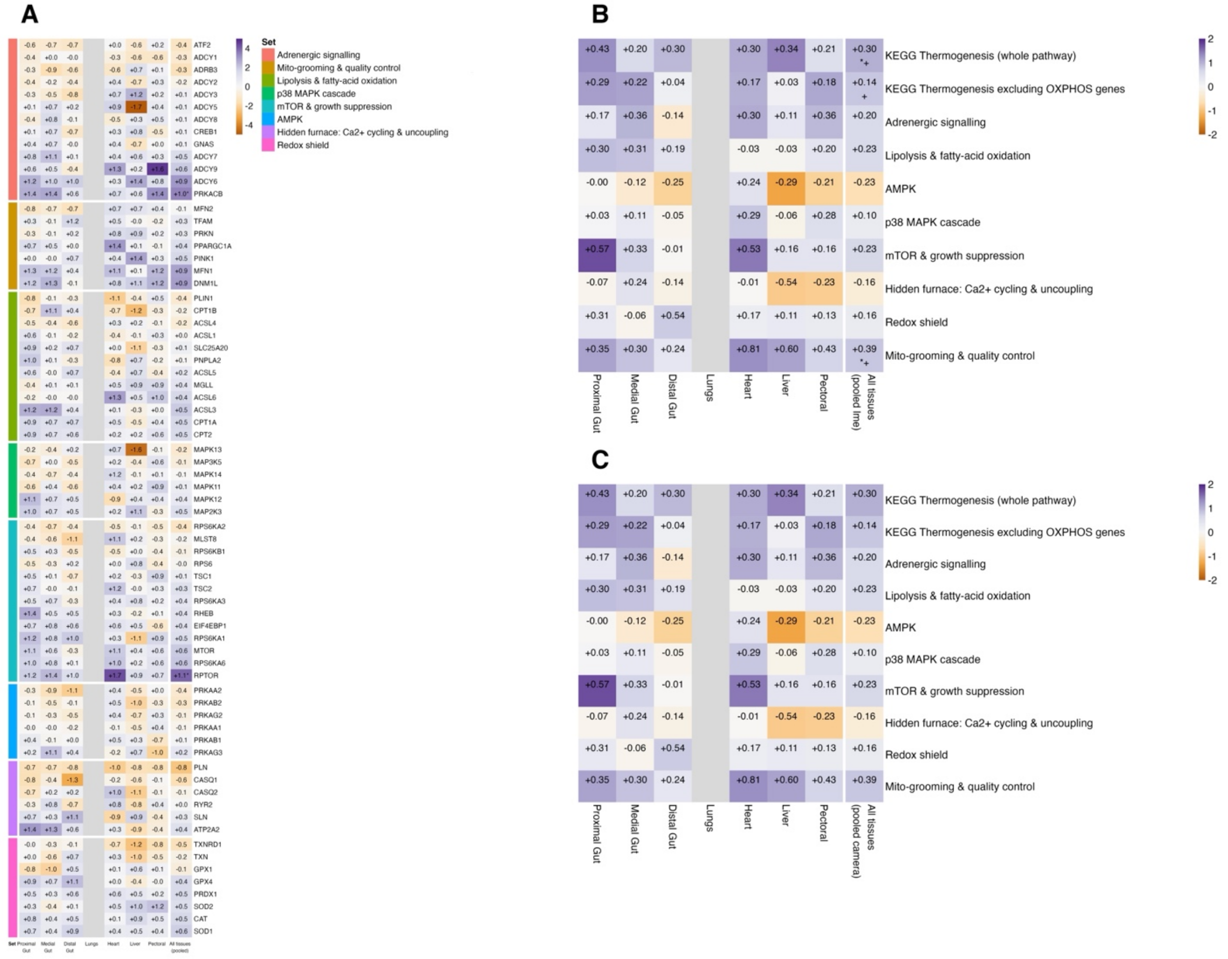


Figure S13: Thermogenesis-related gene sets show modest changes in torpor, led by mitochondrial dynamics and quality control. (A) Gene-level responses for genes in the tested sets, grouped by functional set. Colour shows the limma moderated t for torpor versus normothermy (purple, higher in torpor; orange, lower; scaled to ±5). Numbers give the mean shift in per-tissue Z-score units (torpor − normothermy). Asterisks mark p-adj < 0.05 (*) and p-adj < 0.01 (**) (Benjamini–Hochberg), corrected across all gene × tissue tests for the tissue columns and across genes for the last column. (B) Self-contained tests of whether each set shifts at all. Colour shows Hedges' g (torpor − normothermy; scaled to ±2) and numbers show the mean shift of the set in Z units. Asterisks mark p-adj < 0.05 (*) from Welch's t-test per tissue and from a mixed model for the pooled column. A plus sign (+) marks p-adj < 0.05 in the direction-agnostic mroast test. (C) Competitive tests (camera) with the same colour and number scheme as B. Grey cells were not tested (lungs: fewer than three birds in torpor).

### Supplementary Figures and Tables

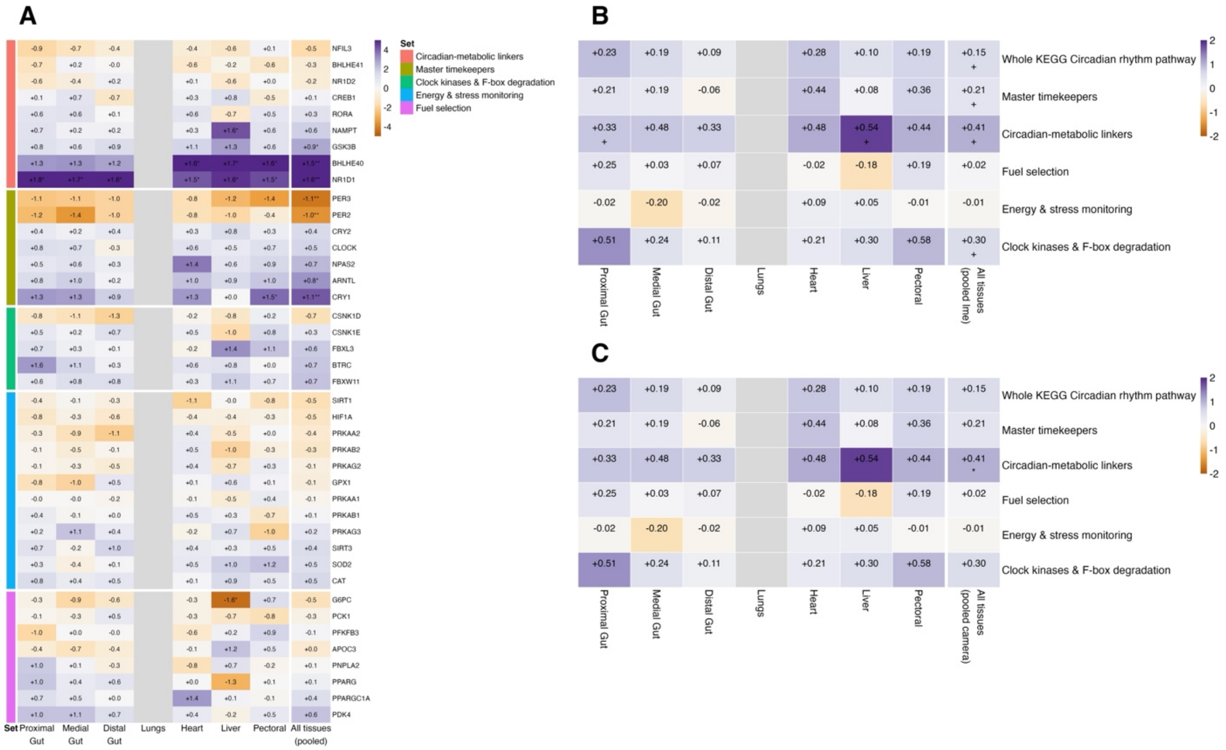


Figure S14: Clock gene sets are rewired in torpor, with opposing shifts among core timekeepers and higher NR1D1 and BHLHE40. (A) Gene-level responses for genes in the tested sets, grouped by functional set. Colour shows the limma moderated t for torpor versus normothermy (purple, higher in torpor; orange, lower; scaled to ±5). Numbers give the mean shift in per-tissue Z-score units (torpor − normothermy). Asterisks mark p-adj < 0.05 (*) and p-adj < 0.01 (**) (Benjamini–Hochberg), corrected across all gene × tissue tests for the tissue columns and across genes for the last column. (B) Self-contained tests of whether each set shifts at all. Colour shows Hedges' g (torpor − normothermy; scaled to ±2) and numbers show the mean shift of the set in Z units. Asterisks mark p-adj < 0.05 (*) from Welch's t-test per tissue and from a mixed model for the pooled column. A plus sign (+) marks p-adj < 0.05 in the direction-agnostic mroast test. (C) Competitive tests (camera) with the same colour and number scheme as B. Grey cells were not tested (lungs: fewer than three birds in torpor).

### Supplementary Figures and Tables

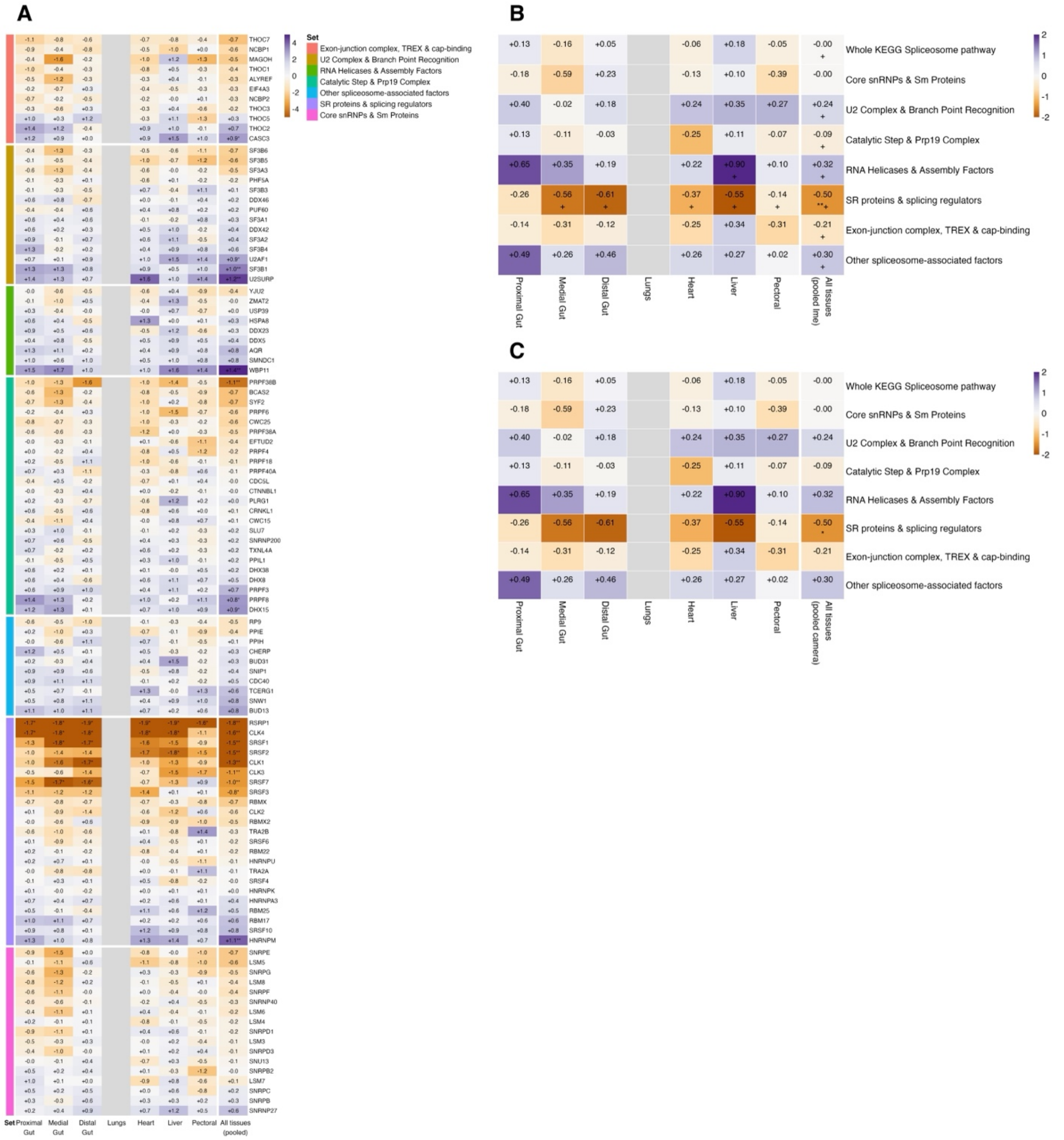


Figure S15: Torpor selectively lowers splicing regulators (SR proteins and CLK kinases), not the spliceosome as a whole. (A) Gene-level responses for genes in the tested sets, grouped by functional set. Colour shows the limma moderated t for torpor versus normothermy (purple, higher in torpor; orange, lower; scaled to ±5). Numbers give the mean shift in per-tissue Z-score units (torpor − normothermy). Asterisks mark p-adj < 0.05 (*) and p-adj < 0.01 (**) (Benjamini–Hochberg), corrected across all gene × tissue tests for the tissue columns and across genes for the last column. (B) Self-contained tests of whether each set shifts at all. Colour shows Hedges' g (torpor − normothermy; scaled to ±2) and numbers show the mean shift of the set in Z units. Asterisks mark p-adj < 0.05 (*) from Welch's t-test per tissue and from a mixed model for the pooled column. A plus

### Supplementary Figures and Tables

sign (+) marks p-adj < 0.05 in the direction-agnostic mroast test. (C) Competitive tests (camera) with the same colour and number scheme as B. Grey cells were not tested (lungs: fewer than three birds in torpor).

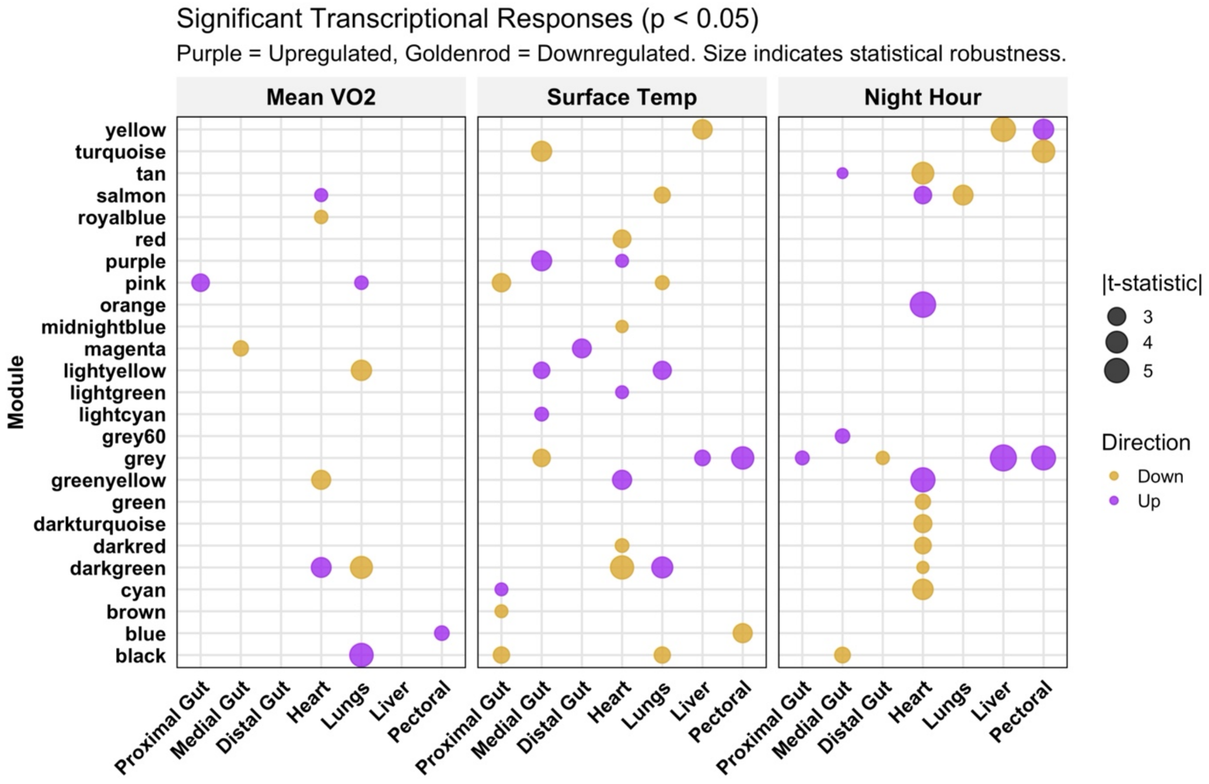


Figure S16: A: Modules are different multiple regression analysis showing correlation of module eigen genes with different physiological traits (Mean $VO_2$, Surface Temperature and Time of the Night). While the colour denotes the directionality of the relationship between the change in expression of module eigengenes and the trait, the size of each bubble denotes the signal-to-noise ratio of the effect (absolute t-statistic). A t-statistic was used to normalize effects of differential scale of change in metabolic rates and surface temperature that characterise entry into torpid states.

### Supplementary Figures and Tables

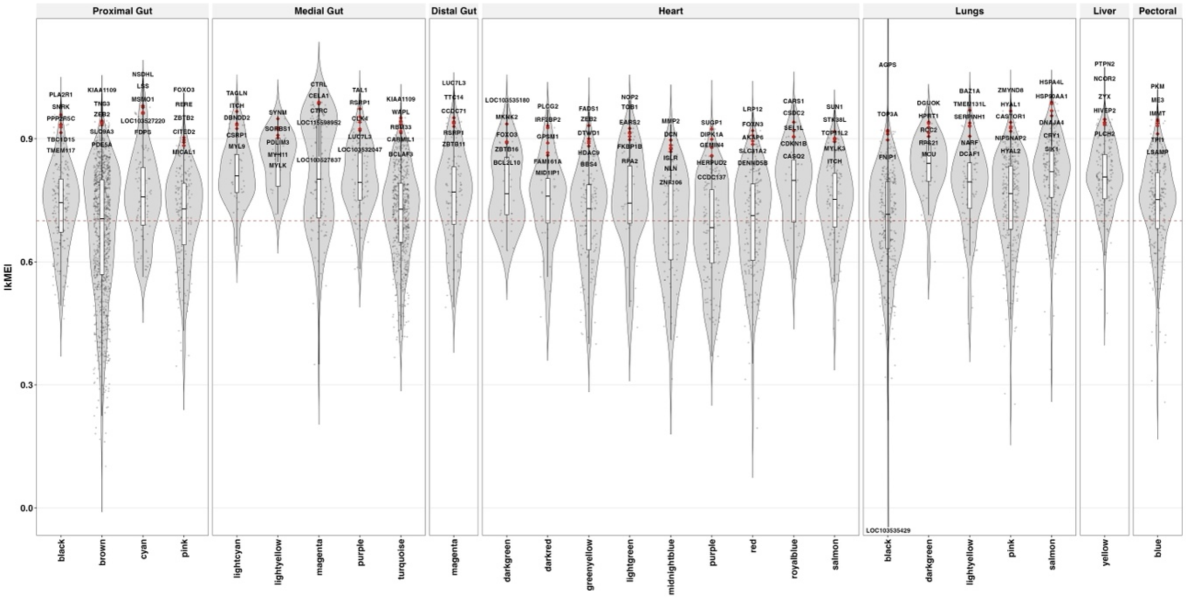


Figure S17: Module membership plot depicting hub genes identified as biologically relevant in prior multiple regression analysis.

### Supplementary Figures and Tables

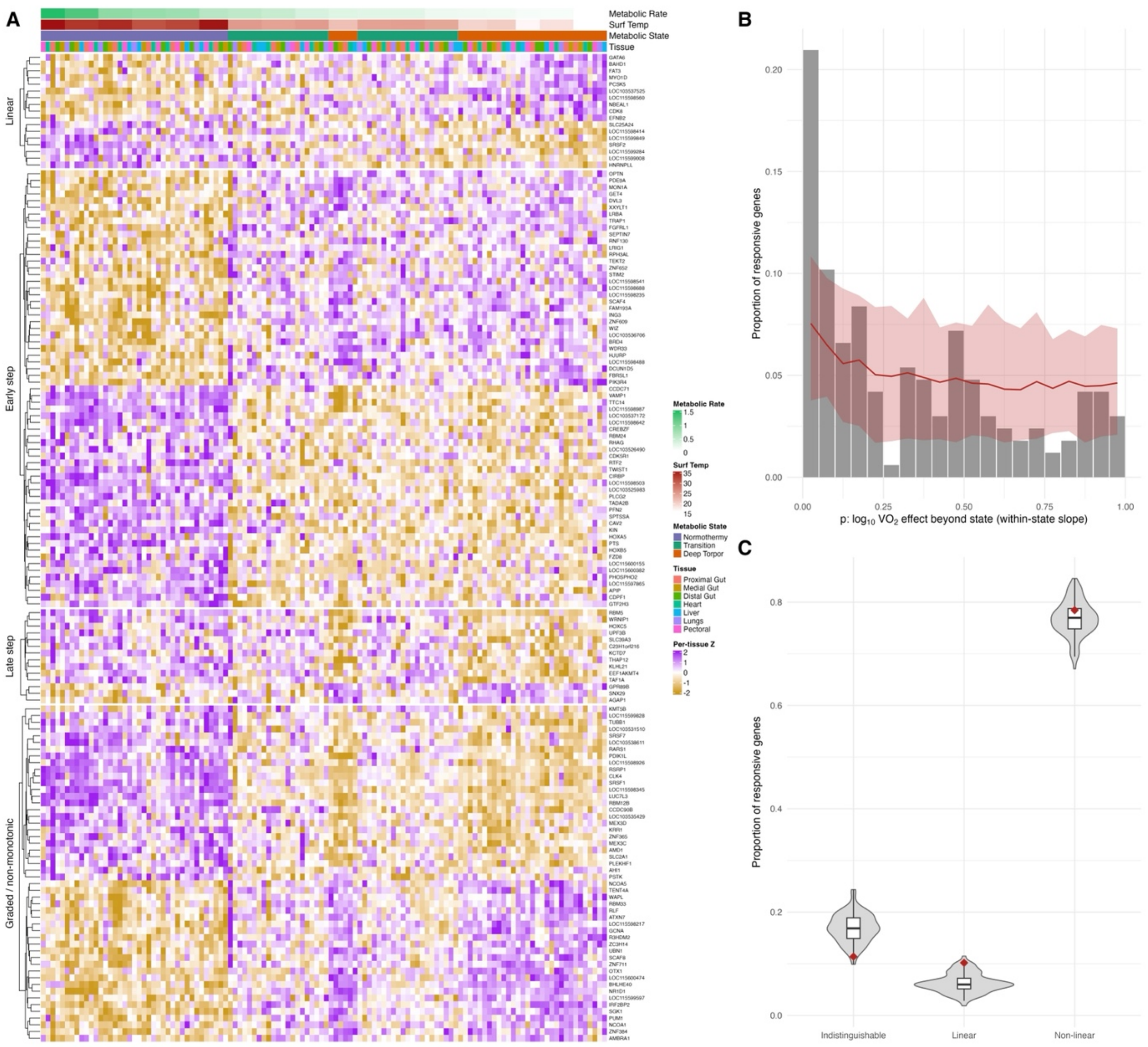


Figure S18: Most $VO_2$-responsive genes differ between states; a smaller set tracks log $VO_2$ continuously. **(A)** Expression of the 148 responsive genes classified as Linear (N = 17) or Non-linear (N = 131; Early step, Late step or Graded/non-monotonic), grouped by class and clustered within class. Each column is one sample (bird × tissue), ordered from highest to lowest metabolic rate. Colours show per-tissue Z-scores of rlog-normalised expression. Top bars show metabolic rate, surface temperature, metabolic state and tissue. Indistinguishable genes (N = 19) are not shown. **(B)** Distribution of p-values for the log $VO_2$ effect beyond state (state-means model vs state-means plus log $VO_2$) across the 167 responsive genes. Grey bars show the observed proportion of genes per bin (bin width 0.05). The red line and band show the mean and 95% range of 100 permutations in which log $VO_2$ was shuffled among birds within state and the full analysis, including gene screening, was repeated. An excess of small p-values over the null indicates a continuous $VO_2$ component (21.0% of genes with $p < 0.05$ vs 7.5% in the null, $p = 0.01$). **(C)** Proportion of responsive genes in each broad class (Linear, Non-linear, Indistinguishable). Violins and boxes show the permutation null (box:

### Supplementary Figures and Tables

median and interquartile range). Red diamonds show the observed proportions. Linear genes exceed the null (10.2% vs 6.3%, $p = 0.02$), whereas Non-linear genes do not (78.4% vs 76.8%, $p = 0.30$). Indistinguishable genes were fewer than in the null (11.4% vs 16.9%). Empirical p = (1 + number of permutations ≥ observed) / (1 + 100).

Table S1: A compilation of the extent of differential gene expression across mammals that express daily torpor vs. hibernation. See PDF attachment.
