## Supplementary Table S1 for "Hummingbird torpor recapitulates key molecular signatures of hibernation without large-scale transcriptomic remodelling"

| Study Citation | Species | Normothermic Temp | Torpid Temp | Ambient Temp | Physio. State | Tissue Type | Measured Unit | Math Fraction | % DE | Notes / Controls |
|---|---|---|---|---|---|---|---|---|---|---|
| **Cubuk et al. (2017a)** | Phodopus sungorus | 36.1 ± 0.7°C | 30.4 ± 0.6°C (sampled during torpor entrance) | 18 ± 1°C | Daily Torpor (entrance phase) | Hypothalamus | Transcripts | 284 / 27,830 | 1.02% | This row is TORPOR ENTRY (entrance phase), not late torpor. Interpret it as an entry-phase contrast. 181 up / 103 down vs ZT-matched normothermic hamsters (n=3 vs 3; $CO_2$ euthanasia). 27,830 = transcripts with RPKM >= 0.1 in at least one sample (Table 2 'Identified genes'). Threshold: fold change >= 1.2 and FDR p <= 0.05 (CLC Genomics Workbench 7.5.1, Exact Test); lenient on fold change. Short-day photoperiod (8:16 L:D) for 12 weeks. Entrance phase, so not comparable with late-torpor values; closest like-for-like comparator to hummingbird entry data. Intro describes torpor as Tb down to ~15C generally, not the sampled animals. Not to be confused with Cubuk, Markowsky & Herwig 2017 PLoS ONE 12:e0186299 (six-gene qPCR). Source: Front. Neurosci. 11:122, doi 10.3389/fnins.2017.00122; Results 'Transcriptomic Expression Analysis' and Table 2. |
| **Deviatiiarov et al. (2021), Commun. Biol. 4:1290** | Mus musculus (C57BL/6J, male, 8.2 weeks) | Not reported in main text | Not reported in main text | Not stated in main text (housed at 21 C before experiments; torpor partly prevented at 28 C, fully at 32 C) | Fasting Torpor (Mid group sampled at ZT-22 on day 2, ~8 h after typical torpor onset at ZT-14; torpor detected from Tb and VO2) | Skeletal muscle (soleus) | Promoters (CAGE clusters) | 866 / 12,862 | 6.73% | Recorded value is the reversible set: 589 up and 277 down promoters (481 + 221 genes) significant (FDR < 0.05) in both Pre to Mid and Mid to Post, the closest analogue to a torpid-vs-baseline contrast. 12,862 = total CAGE peak clusters detected (11,133 clusters associated with 10,617 genes); gene-level cross-check (481 + 221) / 10,617 = 6.61%. Unit is promoters, not genes. Alternatives in the paper: hypometabolic set 330 up + 137 down = 467 (3.63%; torpor vs torpor-prevented HiT and Fed groups); torpor-specific set (intersection of the reversible and hypometabolic sets) 226 up + 61 down = 287 (2.23%; strictest, removes hunger and cold effects). Threshold: edgeR (TMM on TPM), FDR < 0.05, no fold-change cutoff. n: Pre 4, Mid 8, Post 4 (experiment 1); HiT 4, Fed 4; replicate experiment 2 had 2/5/3 animals (38 animals in total). Mid group sampled at ZT-22, ~8 h after typical torpor onset (ZT-14), i.e. established torpor. Fasting-induced torpor in C57BL/6J mice, which enter torpor at TA 12-24 C (paper ref 4); 28 C partly and 32 C fully prevented torpor. Exact TA and Tb of sampled animals are not stated in the main text. Minor internal inconsistencies in the paper: 289 vs 287 torpor-specific promoters; Fig. 1e legend categories sum to 12,863. Source: Commun. Biol. 4:1290. |
| **Hadj-Moussa et al. (2020)** | Microcebus murinus (grey mouse lemur; adult females) | 35-36 C (controls, after arousal from daily torpor) | 30-33 C (minimum Tb during torpor bout) | 24-25 C | Daily Torpor / shallow hibernation (caloric restriction, 5 d at 60% diet; late torpor, sampled at minimum Tb) | Skeletal muscle | miRNA | 4 / 234 | 1.71% (FDR-adjusted p < 0.05) | Recorded value: 4 / 234 miRNAs (miR-2478, miR-1386, miR-4792, miR-8503) = 1.71% at FDR-adjusted p < 0.05. The authors' headline count uses nominal p < 0.01 and fold change > 1.5 (limma eBayes, no multiple-testing correction): 20 / 234 = 8.55% (11 up, 9 down). 234 = conserved mature miRNAs detected in skeletal muscle (small-RNA-seq, mapped to miRBase). Unit is miRNA, not genes. Design: n = 4 torpid vs 4 control adult females, short-day photoperiod, TA 24-25 C; caloric restriction (5 d at 60% of control diet) used to induce torpor. Torpid animals were sampled during a torpor bout at minimum Tb (30-33 C); controls were euthanised after arousal from a daily torpor bout (Tb 35-36 C), not from a never-torpid group. Metabolic rate in torpor is reduced to 20-30% of control levels (70-80% reduction; paper ref 36); torpor is shallow in Tb (~3-6 C) but deep in metabolism, and the paper describes it as shallow daily torpor / multi-day hibernation. Minor inconsistencies in the paper: 30 (Pearson section) and 21 (Fig. 3 legend) DE miRNAs vs 20 in Results and Abstract; mapped reads 58,010 (control) vs 35,957 (torpid). Source: Hadj-Moussa et al. 2020; Results 3.2, Methods 2.1 and 2.6. |

| | | | | | | | | | | |
|---|---|---|---|---|---|---|---|---|---|---|
| **Hrvatin et al. (2020), Nature 583:109-114** | Mus musculus (FosTRAP mice) | >35 C (fed) | 24-35 C (fasting-induced torpor) | 22 C | Fasting Torpor (TRAPed during torpor entry) | Brain (avMLPA neurons; Adcyap1+ neurons) | snRNA-seq (genes; cell-population contrast) | NA / 23,967 | NA | Not usable for the ranges (NA): the paper contains no torpor-vs-normothermic differential-expression contrast. snRNA-seq of the anteroventral preoptic area (avMLPA): 44,669 nuclei from 5 FosTRAP mice (4 TRAPed with 4-OHT as animals entered torpor, 1 non-TRAPed control), used to catalogue cell types (28,103 neurons; 36 neuronal types). The only DE test is TRAPed (n = 139 cells) vs non-TRAPed (n = 5,848 cells) Adcyap1+ neurons within torpid-animal tissue (Seurat FindMarkers, Mann-Whitney U, Bonferroni across the 23,967 genes in the dataset), which identifies markers of the torpor-TRAPed e5 subtype; this is a cell-population contrast, not state remodelling. The result is in Supplementary Table 5; the main text gives no count. 23,967 = genes in the dataset (Bonferroni denominator). Physiology: housed at ~22 C; fed mice Tb > 35.1 +/- 0.2 C; fasting-induced torpor Tb 24-35 C, each bout preceded by a 45.1 +/- 4.6% decrease in metabolic rate; movement down 73.4 +/- 7.0%. Minor inconsistencies in the paper: 342 vs 343 TRAPed neurons; 44,669 nuclei vs 39,562 in the Extended Data Fig. 3 UMAP (after filtering). Source: Nature 583: 109-114; Supplementary Table 5. |
| **Rojas Cuyutupa et al. (2023), Pflugers Arch 475:1149-1160** | Phodopus sungorus (Djungarian hamster) | Not reported (normothermic controls sampled at ZT4) | Not reported (torpor defined as Tb <32 C for >=30 min) | 20 +/- 1 C | Daily Torpor (late torpor, ZT4 torpor nadir) | Whole blood | Genes (de novo transcriptome; human-symbol orthologs) | 11,422 / not reported | NA | Excluded from the ranges (NA): no numerator-denominator pair can be read from the paper. The assembly's gene or contig total and the number of genes tested are not stated. The paper reports 'considerable number of up- or downregulated genes (11,422)' for torpid (HT) vs normothermic (NT) hamsters (DESeq2). The significance threshold for this count is not stated (apparently nominal p <= 0.05, no fold-change cutoff); p <= 0.05 and \|FC\| >= 1.25 were applied afterwards in IPA to select 215 target molecules (148 = 68.8% matched direction in the literature). A genome-annotation total was not used as denominator: the P. sungorus annotation (Psun_UM_1.1; Genome Biol Evol 14:evac080) lists 27,906 genes, which would give 40.9%, but this total is external to the study and the numerator is a nominal-p count. Design: short photoperiod (8 h light); n = 9 hamsters expressed spontaneous daily torpor; on the sampling day 5 torpid (HT) vs 4 normothermic (NT) controls, all sampled at ZT4 (torpor nadir). Torpor defined as Tb < 32 C for >= 30 min (DSI telemetry); Tb of sampled animals not reported. Ambient 20 +/- 1 C. CO2 euthanasia, heart puncture, whole-blood pellet, Illumina HiSeq 2500 2 x 150 bp. Data: GEO GSE239559. Source: Pflugers Arch 475:1149-1160 ('Blood transcriptomics mirror regulatory mechanisms during hibernation - a comparative analysis of the Djungarian hamster with other mammalian species'). |

| | | | | | | | | | | |
|---|---|---|---|---|---|---|---|---|---|---|
| **Faherty et al. (2016), Genome Biol Evol 8:2413-2426** | Cheirogaleus medius (fat-tailed dwarf lemur) | 33.9 +/- 0.1 C (Active 1) / 35.3 +/- 1.8 C (Active 2) (rectal, anaesthetised) | 19.8 +/- 4.1 C (rectal; as low as 15 C) | 25 C (Active) / 15 C (Torpor) | Hibernation (multi-day torpor bout; Cheirogaleus medius is a hibernator; sampled 2-3 days into a bout; Active 1 vs Torpor) | White adipose tissue (tail biopsy) | Genes/transcripts (de novo transcriptome; human-homolog) | 283 / 15,420 (Active 1 vs Torpor) | 1.84% | Recorded value: Active 1 vs Torpor, 283 / 15,420 = 1.84%. Other contrasts: Active 2 vs Torpor 126 / 15,420 = 0.82%; overlap of the two contrasts (genes changing in both) 90 / 15,420 = 0.58%; Active 1 vs Active 2 = 1 gene. Cheirogaleus medius is a hibernator (up to 7 months in the wild); animals were in seasonal multi-day torpor. Denominator: 15,420 transcripts with human homology expressed in at least four samples (Trinity de novo assembly of reads from all 12 samples; 1,295,606 putative transcripts / 776,170 gene models; 158,276 transcripts with a human BLASTX homolog at e < 1e-4; genes with >= 4 samples below 0.5 CPM discarded). The paper uses 'transcripts' and 'genes' interchangeably; after clustering redundant hits it reports 9,540 unique genes with human homology, with which 90 would be 0.94%, 283 2.97% and 126 1.32%. Threshold: edgeR (~condition + RIN), Benjamini-Hochberg FDR < 0.05, no fold-change cutoff; the authors describe these cut-offs as stringent and note that the functional enrichment (none significant) should be interpreted carefully. Design: longitudinal white adipose tissue (tail needle biopsy) from the same animals in Oct (Active 1), Dec (Active 2) and Feb (Torpor); 4 animals sampled (2 M, 2 F, 8-13 yr), 1 removed for weight loss; DE analysis on 3 individuals (n = 3 per state; 12 samples sequenced). Low n and biopsy RNA of variable quality (RIN in the model), so the low percentage reflects limited power as well as biology. State: sampled 2-3 days into a torpor bout (absence of feeding) under anaesthesia (ketamine/midazolam) with rectal Tb; temperatures are group means +/- SD (lowest torpor Tb 15 C). Ta 25 C (Active) / 15 C (Torpor); photoperiod LD 11.5:12.5 / 9.5:14.5 / 10.5: 13.5. Torpor induced by a 66% reduction in caloric input, lowered ambient temperature and shortened days (captive, hibernaculum, torpor Jan-Feb). Source: Genome Biol Evol 8(8):2413-2426, doi 10.1093/gbe/evw163; Methods 'Identification of DE Genes', Tables 1 and 2. |
| **Goropashnaya et al. (2020), Sci Rep 10: 9010** | Urocitellus parryii (arctic ground squirrel) | Not reported for summer-active animals (species ~37 C, literature) | 2.2 +/- 0.3 C (late torpor; 27 +/- 1 C at torpor re-entry) | 2 C (hibernation chamber) | Seasonal Hibernation (late torpor, 80-90% of a bout, 8-12 days) | Skeletal muscle (quadriceps) | Genes | 628 / 12,085 (LT vs SA) | 5.20% | Recorded value: LT vs SA (late torpor vs summer active), 628 DE genes (271 up, 357 down) of 12,085 expressed genes = 5.20%. Other contrasts in Table 1: ET vs SA 383 / 11,956 (3.20%); LT vs ET 173 / 12,042 (1.44%). 1,184 genes (~9.8% of 12,085) are DE in at least one of the three comparisons. Denominator: genes with at least 2 paired reads across all samples in a pairwise comparison (paper: 'expression of 12,000 genes, about 50% of annotated genes'; reads mapped to the 13-lined ground squirrel genome SpeTri2.0, 25,998 genes annotated). Threshold: CLC Genomics Workbench v10, edgeR-style negative-binomial GLM (TMM), Wald test, Benjamini-Hochberg FDR <= 0.05, no fold-change cutoff (observed fold changes start at 1.70). Design: quadriceps, n = 4 per group (LT, ET, SA), adult males except one female in ET; all hibernating animals had completed at least three full-length torpor bouts. LT = 80-90% of bout duration (8-12 days), Tb 2.2 +/- 0.3 C. ET = re-entry into torpor after an interbout arousal (Tb 27 +/- 1 C), i.e. post-arousal entry rather than entry from normothermia. SA = July, after reproductive regression; Tb of SA animals not reported (euthermic interbout arousals reach Tb > 36 C for < 24 h). Hibernating animals held at 2 C with 4:20 L:D (initial housing 20 +/- 2 C, 16:8 L:D). Euthanasia differed by group (torpid animals decapitated without anaesthesia; SA isoflurane; ET sodium pentobarbital); tissue frozen within 9 min of death. Body mass 613.75 +/- 48.69 g (hibernating) vs 719.00 +/- 144.83 g (SA), P = 0.22. Obligate seasonal hibernator (2-3 week bouts, ~98% metabolic depression). Source: Sci Rep 10:9010, doi 10.1038/s41598-020-66030-9; Table 1, Methods 'Animals' and 'Data analysis'. |

| | | | | | | | | | | |
|---|---|---|---|---|---|---|---|---|---|---|
| **Fu et al. (2021), Front Physiol 11:624677** | Ictidomys tridecemlineatus (13-lined ground squirrel) | ~37 C (SpD 35.6-37 C; SA Tb not reported; IBA 30.6-37.4 C) | 5.5-7.2 C (late torpor, LT) | Not stated in main text | Seasonal Hibernation (late torpor LT vs summer active SA; pairwise Wald from Suppl Data Sheets 3-5) | Brain: forebrain, hypothalamus, medulla | Genes (pass-filter, per region) | Late torpor vs SA, padj<=0.001 (Wald): Forebrain 983/11,883; Hypothalamus 2,015/12,184; Medulla 1,367/12,034 (omnibus LRT: 3,266 / 4,998 / 2,781, not used) | 8.27% / 16.54% / 11.36% (padj<0.05: 25.9% / 37.3% / 28.4%) | Recorded values: late torpor (LT) vs summer active (SA), DESeq2 Wald test (Supplementary Data Sheets 3-5, column SA_vs_LT_wald_padj), padj <= 0.001 (the paper's DE threshold): forebrain 983 / 11,883 (8.27%), hypothalamus 2,015 / 12,184 (16.54%), medulla 1,367 / 12,034 (11.36%). At padj < 0.05 the same contrast gives 3,073 (25.9%), 4,546 (37.3%) and 3,416 (28.4%); at strict padj < 0.001 medulla is 1,365 (11.34%). Direction at padj < 0.05: forebrain 1,557 up / 1,516 down; hypothalamus 2,353 / 2,193; medulla 1,957 / 1,459. With an additional \|log2FC\| > 1 filter, 74, 76 and 127 genes remain. Thresholds and denominators reproduce the paper's pairwise counts (e.g. SA vs SpD 2/67/5; Ent vs LT 31/52/28). Denominator: pass-filter genes per region (Fig. 4; rlog >= 7 in at least 4 of 5 individuals in at least one group); 11,882 / 12,184 / 12,033 have a non-missing LT result. The paper's omnibus DESeq2 LRT (padj <= 0.001 across all six states, sex in the model) gives 3,266 / 11,883 (27.5%), 4,998 / 12,184 (41.0%) and 2,781 / 12,034 (23.1%); it is not used because it includes seasonal and interbout-arousal effects. SA is the active-season baseline, so season is confounded with torpor; LT vs entrance phase (Ent) is small (31/52/28 genes), so most of the SA-vs-LT difference is already present on torpor entry. Other numbers: 6,505 hibernation-related DE genes in the abstract (union across regions; 49.9% of 13,035 genes passing the filter in at least one region); 71,149 transcripts (42,881 genes) is the annotation total, not a tested count. Design: n = 5 per state x 6 states x 3 regions = 90 samples. States and Tb: SA (8 Aug); IBA (2-3 h after Tb reached 30 C following > 5 days torpid at < 6.5 C; Tb 30.6-37.4 C); Ent (3-31 Jan; 23-27 C after 10-14 h with Tb > 30 C); LT (31 Jan-26 Feb; Tb 5.5-7.2 C; 80-95% of the previous bout length); Ar (Tb 7-12.8 C); SpD (>= 10 days at Tb 35.6-37 C). SA Tb not given; ambient not stated in the main text (animals as in Hindle & Martin 2013). Source: Front. Physiol. 11: 624677; GEO GSE106947; Fig. 2, Fig. 4, Methods. |
| **Bogren et al. (2017), J Comp Physiol B 187: 735-748** | Ictidomys tridecemlineatus (13-lined ground squirrel) | 37 C (SA and Sp) | 4 C (late torpor) | Not reported in this paper (housing as Grabek et al. 2015) | Seasonal Hibernation (late torpor vs summer active) | Heart; Skeletal muscle (quadriceps); Liver | EDGE-tags (3' tag counts; not genes) | Heart 1,400/30,048; Muscle 950/25,339; Liver 127/25,627 (LT vs SA) | 4.66% / 3.75% / 0.50% | Recorded values: pairwise LT vs SA from Table 2 (DESeq pairwise tests among GLM-significant tags): heart 1,400 / 30,048 (4.66%), skeletal muscle 950 / 25,339 (3.75%), liver 127 / 25,627 (0.50%). These counts are made within the omnibus-significant subset and may undercount true pairwise differences. LT vs Sp: heart 1,288, muscle 428, liver 47. Denominators: tags remaining after filtering (>= 2 TPM in n-1 samples in at least one group), Table 1: 30,048 / 25,339 / 25,627 (starting tags 174,267 / 82,492 / 110,519). Omnibus GLM across all five states (DESeq negative binomial, BH q < 0.05): 5,572 (18.5%) / 11,438 (45.1%) / 4,198 (16.4%); not used because it includes seasonal and IBA effects. Tags within 3 kb of a gene annotation ('DE transcripts'): 3,347 / 6,784 / 2,433; the paper gives no matching gene-level denominator. Unit is EDGE-tags (3' NlaIII tags), not genes. Reference spetri2 (Ensembl release 79); 55.8 / 54.7 / 63.1% of reads uniquely aligned. Design: 74 EDGE-tag libraries; five states per tissue: Sp (spring, Tb ~37 C; n = 5 muscle and liver, n = 4 heart), SA (late July-early August, Tb 37 C, n = 5), IBA (Tb 37 C, 2-3 h after rewarming, n = 5), Ent (Tb 23-27 C, n = 5), LT (80-95% of the prior bout, Tb 4 C, n = 5). Animals exsanguinated under isoflurane, perfused with ice-cold saline, decapitated. Ambient not stated (housing as in Grabek et al. 2015). Obligate hibernator. Source: J Comp Physiol B 187:735-748; Tables 1 and 2, Methods. |

| | | | | | | | | | | |
|---|---|---|---|---|---|---|---|---|---|---|
| **Grabek et al. (2015), eLife 4: e04517** | Ictidomys tridecemlineatus (13-lined ground squirrel) | 37 C (IBA; SA Tb not measured) | 4 C (LT and ET) | Not stated for hibernating groups (spring cold group 4 C) | Seasonal Hibernation (late torpor LT vs summer active SA; re-analysis, not reported in the paper) | Brown adipose tissue (BAT) | EDGE-tags (3' tags); gene-level also given | LT vs SA (our DESeq2 re-analysis of authors' raw counts, padj<0.05): 3,498 / 14,798 tags; 2,795 / 8,089 genes (paper's 3-group omnibus: 2,159 tags / 1,638 genes, not used) | 23.64% (tags) / 34.55% (genes); omnibus 14.59% / 20.25% not used | Recorded values come from our DESeq2 re-analysis (LT vs SA) of the authors' deposited raw counts; they are not reported in the paper. The paper's own test (Suppl. file 3C) is a three-group DESeq test (spring; winter warm = IBA, Ent, SA; winter cold = ET, LT, EAr) and cannot isolate LT vs SA: 2,159 / 14,798 tags (14.59%) and 1,638 / 8,089 genes (20.25%); not used. Re-analysis: pairwise LT (n = 4) vs SA (n = 5), DESeq2 Wald test, Benjamini-Hochberg adjusted p, on the Suppl. file 3A raw EDGE-tag counts restricted to the 14,798 post-filter tags in 3B; technical replicates excluded. Result: 3,498 / 14,798 tags (23.64%) at padj < 0.05 (14,770 tags with a non-missing padj); 2,795 / 8,089 unique Ensembl genes (34.55%) with at least one significant tag. Stricter thresholds: padj < 0.01, 1,798 tags; padj < 0.001, 648 tags. 2,179 tags higher and 1,319 lower in LT; 3,147 tags with an added \|log2FC\| > 1 filter. Caveats: n = 4 vs 5 libraries; LT libraries are larger (~7-14 M counts after filtering vs ~2.5-4 M for SA; handled by DESeq2 size factors); a gene is counted once if any of its tags is significant; SA vs LT is season-confounded; BAT is a thermogenic tissue with unusually large torpor-arousal transcript changes. Denominator: 14,798 EDGE-tags (8,089 unique genes) after normalisation, filtering (>= 2 TPM in n-1 samples in at least one group) and annotation. Unit is EDGE-tags (3' NlaIII tags); tags at different positions within a gene were not combined (possible isoforms), so tag-level and gene-level percentages differ. Tags annotated to the nearest Ensembl (v.71) gene if overlapping or within 3 kb downstream. Design: 38 libraries, 8 sampling groups: ET (Tb 4 C, 5-10% of previous bout, n = 4), LT (Tb 4 C, 80-95% of prior bout, n = 4), EAr (Tb 5-12 C, n = 5), IBA (Tb 37 C, 2-3 h after stabilisation, n = 4), Ent (Tb 23-27 C, n = 5), SA (late July-early August, n = 5; Tb not measured), SpC (n = 5), SpW (n = 6). Ambient for the hibernating groups is not stated in the main text (animals as in Hindle & Martin 2014). Sequencing platform (GAIIx vs HiSeq) included in the model. Euthanasia by exsanguination under isoflurane, perfused with ice-cold saline. Source: eLife 4: e04517, doi 10.7554/eLife.04517; Results 'EDGE-tag library properties', Methods. |
| **Xu et al. (2013), BMC Genomics 14: 567** | Urocitellus parryii (arctic ground squirrel) | Not reported (post-reproductive euthermic control) | Not given for sampled animals (species minimum -2.9 C per Introduction) | Initial housing 20 +/- 2 C; hibernation ambient not stated | Seasonal Hibernation (torpor = early + late torpor pooled; T vs P) | Liver | Genes (custom cDNA array, human-symbol genes) | 916 / 5,314 | 17.24% | Torpor (T) pools early torpor (10-20% of a bout, 4-6 days, n = 4) and late torpor (80-90% of the previous bout, 8-12 days, n = 5); late torpor alone is not separated in the main text. Recorded value: 916 genes DE in T vs post-reproductive euthermic control (P) out of 5,314 unique genes with human symbols = 17.24%. Test: three-state (A arousal, T torpor, P post-reproduction) one-way ANOVA on 9,600 probes, q < 0.05 (Storey-Tibshirani), followed by pairwise Tukey HSD p < 0.05 for each pair; TP signature = 916 genes (AP 823, TA 383). No mean fold-change cutoff for this comparison. A = early arousal (1-2 h after Tb > 30 C, n = 4) + late arousal (7-8 h, n = 4); P = 7 post-reproductive euthermic animals (completed reproductive regression, in moult). Array-based (custom cDNA membrane array, 33P-dCTP, designed from EST clusters), so coverage is partial and biased toward annotated, expressed genes. Denominator: 9,600 clones printed; 6,403 annotated genes, of which 5,314 unique genes with human symbols (redundant probes collapsed to the one with the minimum ANOVA p) were analysed (916 / 6,403 = 14.3%; 916 / 9,600 = 9.5%). Tb of sampled animals not given (species minimum -2.9 C per the Introduction); stage defined by telemetry. Initial housing 20 +/- 2 C, 16:8 L:D; hibernating animals then at 4:20 L:D (ambient during hibernation not stated). Animals trapped near Toolik Lake, Alaska; torpid animals euthanised by decapitation without anaesthesia, aroused and post-reproductive animals under isoflurane; food withheld during hibernation. Source: BMC Genomics 14:567, doi 10.1186/1471-2164-14-567; Methods 'Animals' and 'AGS cDNA microarray analysis'; Results 'Identification of molecular signatures of hibernation'. |

| | | | | | | | | | | |
|---|---|---|---|---|---|---|---|---|---|---|
| **Li et al. (2023), Virulence** | Rhinolophus ferrumequinum (greater horseshoe bat) | Not reported | Not reported | Not reported | Hibernation season (late-hibernation April vs pre-hibernation October; torpor at sampling not confirmed) | Wing tissue (plagiopatagium punch biopsy) | Genes | 2,361 / 20,886 (late vs pre; denominator = annotation total) | 11.30% | Recorded value: late-hibernation (April) vs pre-hibernation (October), 2,361 DEGs / 20,886 = 11.30%. Denominator 20,886 = KEGG total for R. ferrumequinum (organism code rfq): 19,651 protein genes + 1,235 RNA genes (RefSeq / mRhiFer1 annotation). The paper mapped reads to Ensembl release 100 and does not state how many genes were tested, so this is an annotation total from a different annotation. All six pairwise DESeq2 comparisons: 913 (early vs pre), 876 (late vs early), 4,790 (post vs late), 2,361 (late vs pre), 5,379 (post vs pre), 5,682 (post vs early); 8,778 unique DEGs across all comparisons. Only late vs pre is used (closest to late torpor vs baseline). Threshold: DESeq2, Benjamini-Hochberg adjusted P < 0.05 and \|log2FC\| > 1; this fold-change filter makes counts conservative relative to studies without one. Torpor at sampling is not demonstrated: the paper reports no body temperature and does not state that torpor was induced; stages are defined by month. Methods state that active bats were netted as they flew out of the cave at night and hibernating bats were captured from roosts, without specifying which stage was collected by which method. Design: wing tissue (plagiopatagium punch biopsy) from 30 adult bats; pre-hibernation (October, n = 5), early (December, n = 5), late (April, n = 12), post-hibernation (May, n = 8); Oct 2020-May 2021, Jilin Province, NE China. Pd (fungal) infection in 6 of the late-hibernation bats (infected vs uninfected: 3 DEGs). Tb and ambient temperature not reported. Source: Virulence 14:2156185 (2023); Methods 'Sample collection' and 'Differential expression analysis'; Results, Fig. 1c. |
| **Hampton et al. (2013), PLoS ONE 8:e85157** | Ictidomys tridecemlineatus (13-lined ground squirrel) | 35-37 C (active and IBA; rectal) | 6-8 C (torpor; rectal) | 5-7 C (hibernation chamber); 23 C (summer) | Seasonal Hibernation (torpor vs April active; pairwise flags from Suppl Table S2) | Brown adipose tissue (BAT) | Genes (contigs mapped to human mRNAs) | TOR vs APR: 1,533 / 14,573 (authors' pairwise flags in Suppl Table S2). TOR vs OCT (476) and omnibus 2,083 not used | 10.52% (TOR vs APR); TOR vs OCT 3.27% and omnibus 14.29% not used | Recorded value: TOR vs APR (torpor vs April active, post-hibernation), 1,533 / 14,573 = 10.52% (645 higher, 888 lower in torpor), from the authors' pairwise flags in Suppl. Table S2 (2,083 genes x Boolean columns, e.g. 'BAT Tor vs Apr'). April is used as the active baseline (as for Vermillion et al. 2015): April animals were collected in the second and third weeks of April after the chamber was returned to 23 C and represent the post-hibernation state; they have just emerged from a winter of fasting and torpor-arousal cycles, so recovery and reproductive-season effects are possible. Other pairs: TOR vs OCT (October active, pre-hibernation) 476 (3.27%; 245 higher, 231 lower); TOR vs IBA 104 (0.71%); APR vs OCT 882 (6.05%); APR vs IBA 1,215 (8.34%); OCT vs IBA 348 (2.39%). The Table S2 legend describes only the omnibus filter and does not define the pairwise flags. Every flagged pair has a >= 1.5-fold difference in upper-quartile-normalised means, but some genes with >= 1.5-fold differences are not flagged (79 for TOR vs OCT, 57 for TOR vs APR), so an additional pairwise criterion appears to have been applied. Pairwise flags exist only for the 2,083 omnibus-significant genes, so the counts are lower bounds. Omnibus test: DESeq across four collection points (October active, torpor, IBA, April active), prefiltered to genes with a mean >= 100 normalised counts at one point and >= 50% change between any two points (2,631 candidates); BH FDR 0.05 gave 2,083 = 14.29% of 14,573; not used because it includes seasonal and IBA effects. Denominator: 14,573 distinct transcripts mapped to human RefSeq mRNAs (BLASTn) from Trinity contigs (BAT reads plus skeletal muscle, heart, hypothalamus and cortex reads); not a count of genes tested after expression filtering. Design: 12 male + 12 female 13-lined ground squirrels; 6 animals per collection point; RNA from one male and one female pooled, giving n = 3 libraries per point (12 in total; 179,979,787 BAT reads). Mitochondrial-encoded reads ~40% of BAT reads. Torpor sampled in January; phase of the bout not stated. Rectal Tb: torpor 6-8 C, active and IBA 35-37 C. Hibernation chamber 5-7 C in constant darkness (ambient lowered to 5 C on 1 November, raised to 23 C in March/April); summer room 23 C, 12:12 L:D. Animals killed by decapitation under 5% isoflurane between 10:00 and 15:00. Upper-quartile normalisation, DESeq. Source: PLoS ONE 8(12): e85157, doi 10.1371/journal.pone.0085157; Results and Methods. |

| | | | | | | | | | | |
|---|---|---|---|---|---|---|---|---|---|---|
| **Vermillion et al. (2015), Physiol Genomics 47: 58-74** | Ictidomys tridecemlineatus (13-lined ground squirrel) | 35-37 C (OCT and IBA; rectal) | 6-8 C (torpor; rectal) | 5-7 C (hibernation chamber); 23 C (October/April) | Seasonal Hibernation (torpor vs April active, post-hibernation; chosen baseline) | Heart; Skeletal muscle (quadriceps) | Genes (>=10 reads, human-symbol) | Heart 967/8,285; Muscle 1,317/8,278 (TOR vs APR). TOR vs OCT (not used): 786 and 938 | 11.67% / 15.91% (TOR vs APR); TOR vs OCT 9.49% / 11.33% not used | Recorded values: TOR vs APR (torpor vs April active, post-hibernation): heart 967 / 8,285 (11.67%), skeletal muscle 1,317 / 8,278 (15.91%). April animals were collected in the second and third weeks of April after removal from the hibernation chamber at the end of March; the authors state that April reflects the post-hibernation state, and use it as the reference. These animals have just emerged from a winter of fasting and torpor-arousal cycles, so recovery and reproductive-season effects are possible. TOR vs OCT (October active, pre-hibernation): heart 786 (9.49%), muscle 938 (11.33%). Both sets from Table 2 (post hoc DESeq on omnibus-DE genes; prefilter >= 50% change between the two points; BH FDR 0.05). Other pairs, heart / muscle: OCT-APR 296 / 524; OCT-IBA 483 / 688; TOR-IBA 182 / 109; IBA-APR 749 / 1,085. Omnibus DE (analysis of deviance across APR, OCT, TOR, IBA; >= 50% change between any two points and mean >= 100 reads at one point; BH FDR 0.05): 1,076 heart (12.99%) and 1,466 muscle (17.71%); not used because it includes seasonal and arousal effects. Pairwise counts are taken only among omnibus-significant genes and may undercount true pairwise differences; both tests have an abundance / fold-change prefilter. October animals 'are experiencing shallow torpor bouts' (Tb as low as 20 C for up to 24 h) although active at death. Denominators: genes with >= 10 reads in total across the four time points that map to protein-coding human-symbol genes: 8,285 (heart), 8,278 (muscle), 8,229 shared (the contig assembly gave 14,332 and 14,169 distinct transcripts). Design: 24 thirteen-lined ground squirrels (12 M, 12 F); 3 M + 3 F per collection point, RNA pooled into three samples per point (each pooled across sexes), n = 3 libraries per point per tissue (12 per tissue). TOR: >= 3 days in a torpor bout, January-February, no visible signs of arousal, rectal Tb 6-8 C. IBA: aroused naturally, active, Tb 35-37 C. OCT: first 2 weeks of October. APR animals held at 12:12 L:D, 23 C, food ad libitum after removal from the chamber. Hibernation chamber: constant darkness, 5-7 C, no food. Killed by decapitation under isoflurane; tissue frozen < 10 min. DESeq v1.6.1, upper-quartile normalisation. Heart: both atria and ventricle (sagittal half); skeletal muscle: quadriceps femoris. Source: Physiol Genomics 47:58-74, doi 10.1152/physiolgenomics.00108.2014; Methods and Results, Table 2, Supplemental Tables S1-S2. |

| | | | | | | | | | | |
|---|---|---|---|---|---|---|---|---|---|---|
| **Gillen et al. (2021), Front Physiol 12: 662132** | Ictidomys tridecemlineatus (13-lined ground squirrel) | ~37 C (SpD 35.5-37 C; IBA 30.6-37.7 C; SA Tb not given) | 5.5-5.7 C (LT, GRO-seq group); 7.9-12.8 C (Ar, RNA-seq) | Not stated in main text | Seasonal Hibernation (GRO-seq nascent transcription: late torpor LT vs SA; RNA-seq has no torpor group) | Liver | Genes (GRO-seq pass-filter, nascent transcription) | GRO-seq LT vs SA: 1,496 / 4,705 (padj<0.001). RNA-seq has no torpor group (omnibus 3,120/10,370 not used) | 31.80% (GRO-seq, padj<0.001); 42.04% at padj<0.01; 53.13% at padj<0.05. RNA-seq omnibus 30.1% not used | Recorded value: GRO-seq (nascent transcription), late torpor (LT) vs summer active (SA), DESeq2 Wald test (Supplementary Data Sheet 4, column padj-SA_vs_LT): 1,496 / 4,705 pass-filter genes (31.80%) at padj < 0.001, the paper's DE threshold; 1,978 (42.04%) at padj < 0.01; 2,500 (53.13%) at padj < 0.05 (1,182 higher, 1,318 lower in LT); 1,277 with \|log2FC\| > 1 and padj < 0.05. 4,703 genes have a non-missing padj. GRO-seq measures engaged polymerase (nascent transcription), not steady-state mRNA, so it is a different quantity from the mRNA-level contrasts in other rows. Small, uneven n: n = 3 for SA, IBA, Ent and ET but n = 2 for LT (LT71 removed: poor quality and PCA outlier). The denominator is a GRO-seq pass-filter set (>= 25 TPM in SA or one hibernation group; transcription units redefined by Fstitch), which favours highly transcribed genes and is smaller than the RNA-seq set (10,370), so the percentage is inflated relative to whole-transcriptome denominators. The RNA-seq dataset (steady-state RNA) has no torpor group: its five states are SA, IBA, Ent, Ar (late arousal, Tb 7.9-12.8 C, rising) and SpD, n = 5 each. Its omnibus 3,120 / 10,370 (30.1%; DESeq2 LRT, padj <= 0.001, sex in the model) is therefore not a torpor contrast and is not used; the closest RNA-seq contrast, SA vs Ar (arousal), is 1,708 / 10,370 (16.47%) at padj < 0.001 (37.3% at padj < 0.05). GRO-seq omnibus LRT (padj <= 0.001): 2,417 / 4,705 = 51.4% (the paper's text states 56%). Other GRO-seq pairs at padj < 0.001: ET vs LT 33; SA vs ET 1,726; LT vs IBA 1,515; LT vs Ent 679; IBA vs SA 175. Only 770 genes were DE in both datasets. Design: liver. States and Tb: SA (8 Aug); IBA (2-3 h after Tb reached 30 C following > 5 days torpid at < 6.5 C; Tb 30.6-37.7 C; 11 Feb-26 Mar); Ent (3-31 Jan; 23-27 C after 10-14 h with Tb > 30 C); Ar (28 Dec-22 Jan; Tb 7.9-12.8 C); SpD (21 Mar-30 Apr; >= 10 days at Tb 35.5-37 C); ET (3 Dec-13 Jan; Tb < 30 C after an IBA, 8-16% of the previous bout); LT (31 Jan-22 Feb; Tb 5.5-5.7 C; 80-95% of the previous bout). Ambient not stated in the main text. Source: Front. Physiol. 12:662132, doi 10.3389/fphys.2021.662132; Results 'RNA-Seq Data Analysis', Figs 2C and 5B, Methods. |
| **Lei et al. (2014), PLoS ONE 9:e107746** | Rhinolophus ferrumequinum (greater horseshoe bat) | ~25 C (surface temperature of active bats; core Tb not reported) | ~5 C (surface temperature of torpid bats; core Tb not reported) | Not reported | Seasonal Hibernation (winter torpid vs summer active) | Brain (whole, without cerebellum) | Genes (de novo assembly, non-redundant contigs) | NA (one library per state: no biological replication) | NA | Percentage set to NA: there is one library per state (SRA accessions SRR1048140, active; SRR1048142, torpid), so there is no biological replication. For reference, the authors report 1,573 DE genes (1,038 up, 535 down in torpor; edgeR, Benjamini-Hochberg FDR < 0.05 and fold change > 2) out of 11,214 non-redundant contigs (14.03%); not used in the ranges. Denominator: contigs remaining after CD-HIT (95% identity; 11,279 contigs removed) from a Trinity de novo assembly of reads from both states; 9,910 (93% of the annotated ones) were assigned GO terms. This is an assembly-based gene set, not a count of genes tested, and no reference genome was used. The 'replicates' are two sequencing lanes per state (Table 1: Active 1/2 and Torpor 1/2; 22.5 M / 17.5 M and 12.8 M / 17.3 M used reads); the paper does not state whether RNA was pooled. edgeR dispersion would therefore reflect technical rather than biological variance, and the count may be inflated. Design: six male summer-active bats (24 June 2009) and six male winter-torpid bats (11 December 2009), wild-caught in Fish cave, Anhui Province, China; whole brain without cerebellum; Illumina GA II, 75-bp paired-end. Temperatures are surface temperatures (~25 C active, ~5 C torpid); core Tb and ambient temperature not reported. Torpor phase not stated (single December collection in the cave). Bats euthanised by respiratory hyperanaesthesia followed by thoracotomy; tissue in RNAlater. Source: PLoS ONE 9(9):e107746, doi 10.1371/journal.pone.0107746; Results 'De novo assembly and functional annotation' and 'Differentially expressed genes'; Supplementary Table S2. |

<table>
<tr><th></th><th></th><th></th><th></th><th></th><th></th><th></th><th></th><th></th><th></th><th></th></tr>
<tr><td>Srivastava et al. (2019), DNA Research 26: 37-44</td><td>Ursus americanus (American black bear)</td><td>Not reported (37 C is a species literature value)</td><td>Not reported (29 C is a species literature value)</td><td>Not reported</td><td>Seasonal Hibernation (late fall vs early spring; neither group torpid)</td><td>Kidney</td><td>Genes (protein-coding, annotation total)</td><td>169 / 18,091</td><td>0.93%</td><td>Not a torpor contrast: late-fall (just before hibernation) vs early-spring (~1-2 weeks after emergence) kidneys; neither group was torpid. Documented exclusion; not used in the ranges. Recorded value: 169 protein-coding genes DE between spring and fall (101 down, 68 up after hibernation; DESeq2, adjusted P / FDR <= 0.05, no fold-change cutoff; fold changes 1.8-fold down to 2.4-fold up; only CISH, SLCO1C1 and NDST3 exceeded 2-fold) / 18,091 = 0.93% (0.57% if all 29,624 annotated genes were used). Denominator: coding genes in the CAT-annotated black bear assembly (29,624 genes in total; 22,081 protein-coding transcripts; annotation projected from dog Ensembl v87 and other species); the paper does not state how many genes were tested. Body and ambient temperatures are not reported; the temperatures in the table are species literature values. Design: wild Maine bears sampled by hunters; fall (JAX118, 134, 141, 146, 160, 165) and spring (JAX101, 103, 106, 113, 136, 140), three males and three females per time point; spring sample 101 (male) removed as a PCA outlier, leaving 11 bears (6 fall, 5 spring); batch effect corrected; ~60 M reads per sample (Illumina TruSeq Stranded Total RNA with Ribo-Zero Gold). Source: DNA Research 26(1):37-44 (2019), doi 10.1093/dnares/dsy036; Results 'Differential expression between fall and spring', Sections 2.1, 2.4, 3.2.</td></tr>
<tr><td>Zhang et al. (2024), Front Neurosci 18: 1501223</td><td>Tamias sibiricus (Siberian chipmunk)</td><td>Not reported (surface Ts ~35 C in ACT, read from Fig 1B)</td><td>Not reported (surface Ts ~23 C E-DS; ~13 C M-DS, read from Fig 1B)</td><td>10 +/- 2 C (hibernation experiment)</td><td>Seasonal Hibernation (late torpor E-DS vs pre-hibernation ACT)</td><td>Hypothalamus</td><td>Genes (Corset clusters from de novo Trinity assembly)</td><td>254 / not reported</td><td>NA (denominator not reported)</td><td>Percentage left as NA: the number of genes or clusters tested is not reported in the paper or its supplement. Reference-free design (Trinity de novo assembly; transcripts clustered into 'genes' with Corset); the only unigene counts given are Swiss-Prot hits (Homo sapiens 14,488; Mus musculus 7,545; Bos taurus 3,000), which are not a total. Methods describe both a reference-independent assembly (2.3) and HISAT2 genome alignment (2.6), and the splicing-event table uses chromosome-based gene IDs, so the denominator would depend on an annotation (the T. sibiricus genome, PRJNA1192718, is under review) that is not provided. Pairwise DEG counts (Fig. 1G; text): ACT vs M-DS 100; ACT vs E-DS 254; ACT vs M-EUT 477; ACT vs E-EUT 19; M-DS vs M-EUT 472; E-DS vs E-EUT 122. Late-torpor contrast recorded: ACT vs E-DS = 254 (E-DS = end of hibernation, sampled 1-2 h after > 36 h of continuous torpor, end of March, at the last re-entry into torpor); ACT = pre-hibernation active animals sampled in November. Threshold: DESeq2, adjusted p < 0.05 'and a threshold fold change' (Methods 2.5 gives no value; the volcano plots state |logFC| > 1.5); it is not stated whether the cutoff was applied to the Fig. 1G counts. Design: n = 3 per group, 15 animals, five groups (ACT, M-DS, M-EUT, E-DS, E-EUT), hypothalamus; bulk RNA-seq, Illumina NovaSeq 6000; 15 libraries, 26.8-32.6 M read pairs each, 85.0-88.5% mapped. Temperatures: no core Tb; surface temperature (Ts) from thermal imaging, read approximately from Fig. 1B: ACT ~35, M-DS ~13, M-EUT ~28, E-DS ~23, E-EUT ~35 C (only ACT vs M-DS tested, p < 0.01). Ambient 10 +/- 2 C (40 +/- 10% humidity, 24 h dark) during the hibernation experiment; before that animals were kept at 22 C, 12: 12 L:D, with a 2-month fattening period. Caveats: ACT animals are pre-hibernation (November), so the contrast mixes torpor with season, photoperiod and fattening effects; E-DS Ts (~23 C) is much higher than M-DS (~13 C) and the only criterion for E-DS is > 36 h of torpor; ACT vs E-EUT is only 19 DEGs, so the 254 reflects the torpid state rather than season. Data: BioProject PRJNA1101579. Source: Front. Neurosci. 18:1501223 (2024).</td></tr>
</table>
